# NLRP11 promotes non-canonical inflammasome activation in human macrophages by enhancing caspase-4 recognition of cytosolic lipopolysaccharide

**DOI:** 10.64898/2026.02.02.703338

**Authors:** María Luisa Gil-Marqués, Pascal Devant, Jonathan C. Kagan, Marcia B. Goldberg

**Affiliations:** Division of Infectious Diseases, Department of Medicine, Massachusetts General Hospital, Boston, Massachusetts, USA; Department of Microbiology, Blavatnik Institute, Harvard Medical School, Boston, Massachusetts, USA; Broad Institute, Cambridge, Massachusetts, USA; Division of Gastroenterology, Boston Children’s Hospital and Harvard Medical School, Boston, MA, USA

**Author notes:** Address correspondence to María Luisa Gil-Marqués. Present address: Gladstone-UCSF Institute of Genomic Immunology, San Francisco, CA, USA.

**Keywords:** NLRP11, Caspase-4, Lipopolysaccharide (LPS), Inflammasome, Gasdermin-D, *Shigella flexneri*, Macrophage

## Abstract

Innate immune detection of Gram-negative bacteria depends on sensing of cytosolic lipopolysaccharide (cLPS) by the non-canonical inflammasome, mediated in humans by NLRP11 and caspase-4 (CASP4). Activation of this pathway in human macrophages triggers gasdermin-D activation and pyroptotic cell death. Although CASP4 directly binds LPS in vitro, additional host factors are required for efficient activation in vivo. Here, we show that NLRP11, a primate-specific pattern recognition receptor, facilitates CASP4 recognition of cLPS and promotes non-canonical inflammasome activation. NLRP11 functions upstream CASP4, forming an ASC-independent complex that requires a conserved CASP4 p20 residue, binds cLPS, and enhances CASP4-dependent LPS recognition. Mutational analyses demonstrate that in human macrophages, in addition to LPS binding and CASP4 catalytic activity, CASP4 interaction with NLRP11 is essential for efficient pyroptosis. Together, these findings establish NLRP11 as a primate-specific determinant that enhances CASP4-mediated cLPS detection and non-canonical inflammasome activation, revealing a mechanism for human-specific regulation of innate immunity.

## Introduction

During human infection, initial innate immune responses to bacterial pathogens are mounted by macrophages, in which pattern-recognition receptors detect pathogen-associated molecular patterns and initiate signaling programs that promote pathogen clearance [1]. Innate immune detection of Gram-negative bacteria relies largely on recognition of lipopolysaccharide (LPS), a conserved glycolipid component of the bacterial outer membrane [2]. LPS encountered extracellularly or within endosomes is sensed by the Toll-like receptor 4 (TLR4) signaling complexes, whereas LPS that gains access to the cytosol activates a distinct surveillance pathway known as the non-canonical inflammasome [3–5]. Inflammasomes are cytosolic multiprotein complexes that assemble in response to pathogen-associated molecules, inducing a form of cell death called pyroptosis [5]. In human cells, cytosolic LPS (cLPS) is detected by the inflammatory caspases caspase-4 (CASP4) and caspase-5 (CASP5), while mice rely on the CASP4 homolog, caspase-11 (CASP11) [6,7]. Upon LPS engagement, these caspases undergo proximity-induced activation, activating the pore-forming protein gasdermin D (GSDMD) by proteolytic processing and triggering pyroptotic cell death [7,8]. This process promotes secondary activation of the canonical NLRP3 inflammasome, resulting in amplified inflammatory signaling [9–11].

Although direct binding of LPS to CASP4 or CASP11 triggers caspase oligomerization and activation in vitro [5,7], accumulating evidence suggests that additional host factors influence the efficiency, specificity, and magnitude of cLPS sensing in intact cells [12–14]. This issue is particularly relevant in humans, where CASP4-dependent responses are more robust and complex than those observed in murine systems [5,15,16], pointing to the existence of additional species-specific regulatory mechanisms. Notably, humans exhibit approximately 10^4^-fold greater sensitivity to LPS-induced inflammation and sepsis than mice [17,18], yet because of their ease as a model system, mice are widely used in studies of sepsis and inflammasome signaling, leading to important gaps in our understanding of how cLPS is detected and processed in human cells.

A key regulator of human-specific cLPS sensing is NLRP11, a primate-restricted member of the NOD-like receptor (NLR) family, that is absent in mice [11,16]. The role of NLRP11 in non-canonical inflammasomes was initially identified in a genetic screen in a human myeloid-derived cell line as a host factor required for cell death during infection with the intracellular Gram-negative pathogen *Shigella flexneri* [16,19]. NLRP11 interacts with a subset of inflammatory caspases, binds cLPS, and promotes CASP4-dependent responses to intracellular Gram-negative bacteria in human macrophages. Loss of NLRP11 impairs pyroptosis induced by Gram-negative bacterial infection or LPS electroporation, indicating its essential role in CASP4 inflammasome activation [9,16]. Despite these findings, the molecular basis of NLRP11 function in non-canonical inflammasome signaling remains unclear.

Here, we define the mechanistic role of NLRP11 in non-canonical inflammasome activation during initial contact with pathogens in human macrophages. We show that NLRP11 acts upstream of CASP4 to promote cLPS sensing, CASP4 activation, GSDMD cleavage, and pyroptotic cell death. We demonstrate that NLRP11 enhances CASP4-dependent LPS binding. Together, our findings establish NLRP11 as a primate-specific regulator of the non-canonical inflammasome that scaffolds CASP4 to optimize detection of intracellular LPS.

## Results

### CASP4 is downstream of NLRP11 in pyroptotic cell death induced by cytosolic lipopolysaccharide

To extend our prior findings that the CASP4 inflammasome activation is defective in human-derived THP-1 macrophages lacking *NLRP11* [16], we tested whether NLRP11 and CASP4 are epistatic with respect to the non-canonical inflammasome cell death pathway. We compared cell death triggered by cytosolic *S. flexneri* or LPS in human THP-1 macrophages containing deletions in NLRP11, CASP4, or both. To better represent the initial contact of *S. flexneri* with macrophages, these experiments, and all experiments described in this paper, were carried out without interferon priming. In THP-1 macrophages lacking *NLRP11*, *CASP4*, or both *NLRP11* and *CASP4*, cell death measured by lactate dehydrogenase (LDH) release during *S. flexneri* infection was reduced by more than 50% at 5 h 40 min of infection (p<0.0001) (Figure 1A). No significant differences were observed between *NLRP11^−/−^*, *CASP4^−/−^*, or double-knockout cells, indicating that NLRP11 and CASP4 function within the same pathway.

**Figure 1.**
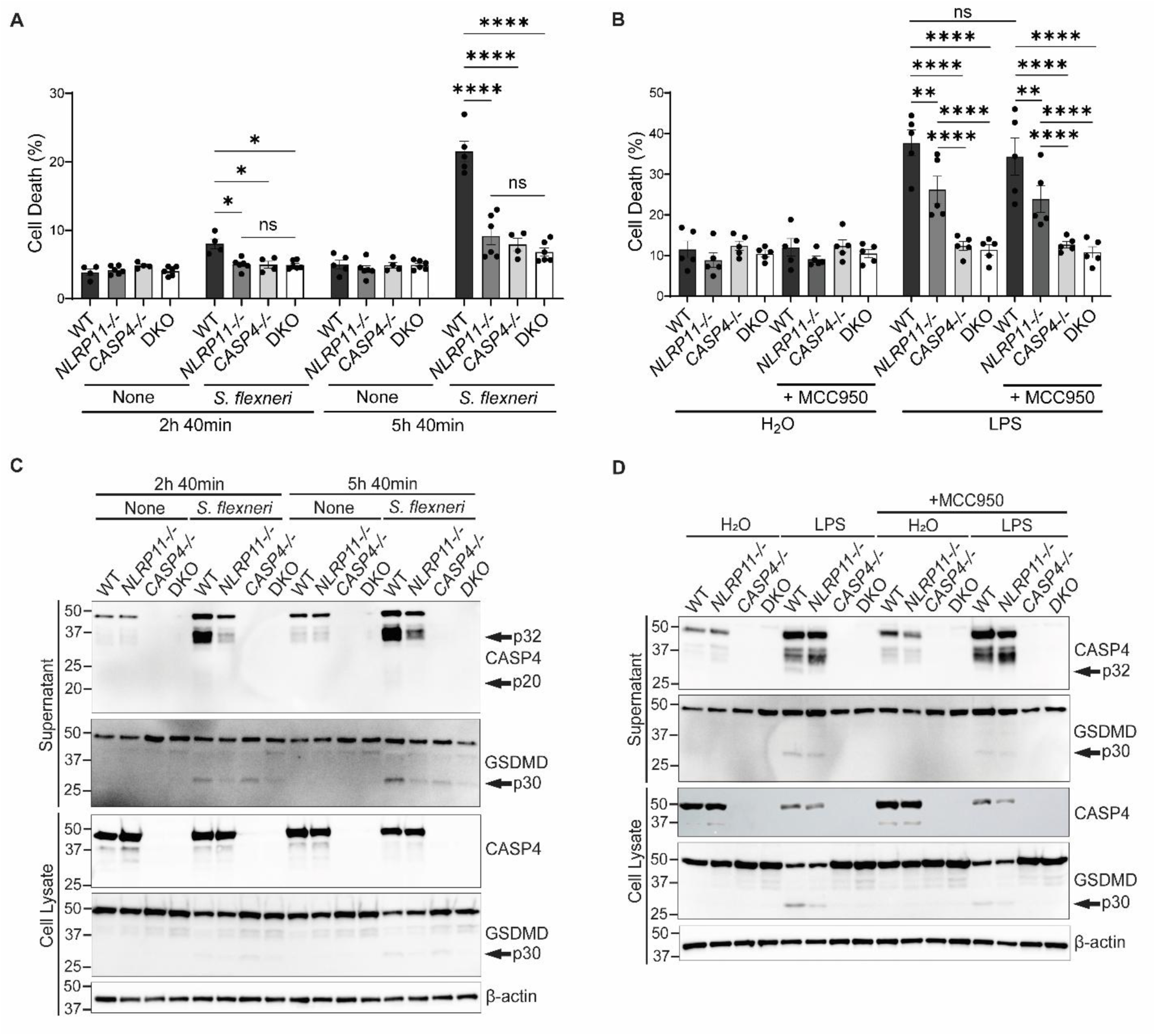
NLRP11 and CASP4 but not NLRP3 are required for efficient cell death of THP-1 macrophages induced by intracellular *S. flexneri* infection or by LPS electroporation. (**A-B**). Cell death of THP-1 macrophages upon infection with *S. flexneri* 2457T (**A**), or electroporation with *S. enterica* serovar Minnesota LPS (**B**), measured by LDH release, as percentage of Triton X-100-induced cell lysis. N ≥ 4 biological replicates. (**C-D**). NLRP11 is required for the processing of CASP4 and gasdermin D (GSDMD) induced by *S. flexneri* infection (**C**), or *S. enterica* LPS electroporation (**D**). CASP4 is required for GSDMD processing induced by *S. flexneri* infection (**C**), or *S. enterica* LPS electroporation (**D**). Processing of CASP4 yields a p32 and/or p20 polypeptide, and processing of GSDMD yields a p30 polypeptide. Representative western blots. Molecular weight markers in kDa. DKO, *NLRP11^−/−^ CASP4^−/−^* double knockout. Mean ± SEM. Two-way ANOVA with Tukey’s post hoc test *, P<0.05; **, P<0.01; ****, P<0.0001.

Introduction of *Salmonella enterica* serovar Minnesota lipopolysaccharide (LPS) or *Escherichia coli* LPS directly into the cytosol by electroporation similarly induced cell death that was dependent on both NLRP11 and CASP4. Following either *S. enterica* or *E. coli* LPS electroporation, LDH release was reduced by approximately 30% in *NLRP11^−/−^* macrophages compared to wildtype (WT) macrophages (p<0.01 and p<0.0001, respectively) and by 65% in *CASP4^−/−^* and *NLRP11^−/−^ CASP4^−/−^* macrophages (p<0.0001) (Figures 1B and S1A). In the case of LPS electroporation, but not *S. flexneri* infection, cell death in *CASP4^−/−^* and *NLRP11^−/−^ CASP4^−/−^*macrophages was decreased compared with *NLRP11^−/−^* macrophages (p<0.0001; Figures 1B and S1A versus 1A), indicating that in response to purified LPS, but not bacterial-associated LPS, CASP4 contributes to cell death through both NLRP11-dependent and NLRP11-independent mechanisms. Pretreatment with the NLRP3 inhibitor MCC950 did not alter cell death in any of these macrophages following LPS electroporation (Figure 1B), indicating that NLRP11- and CASP4-dependent cell death in response to cytosolic LPS occurs independently of NLRP3. Together, these results indicate that CASP4 lies downstream of NLRP11 in an NLRP3-independent macrophage cell death pathway activated by cytosolic bacterial LPS.

### NLRP11 is required for noncanonical inflammasome activation of CASP4 and GSDMD

To further investigate the role of NLRP11 in non-canonical inflammasome signaling, we examined its contribution to CASP4 and GSDMD activation in response to cytosolic *S. flexneri* or LPS. Activation of CASP4, which involves auto-processing to generate the protease-active p32 and p20 domains, was reduced in culture supernatants of *NLRP11^−/−^* macrophages at 5 h 40 min of *S. flexneri* infection by 75% (p<0.0001) and by 45% (p<0.05) and by 40% following *S. enterica* or *E. coli* LPS electroporation, respectively (Figures 1C, 1D and S1B-S1E). Total levels of pro-CASP4 (the combination of that in supernatants and cell lysates) remained unchanged under these conditions, indicating that the NLRP11-dependent increase in released CASP4 active domains was not due to altered protein expression. Similarly, activation of GSDMD, measured by release of its active p30 domain, was diminished by more than 60% in *NLRP11^−/−^* (p<0.001), *CASP4^−/−^* (p<0.0001), and *NLRP11^−/−^ CASP4^−/−^* (p<0.0001) macrophages 5 h 40 min of infection, whereas levels of full-length GSDMD in supernatants and cell lysates were independent of NLRP11 or CASP4 (Figures 1C and S1C). Following *S. enterica* or *E. coli* LPS electroporation, GSDMD processing was reduced by 40% in *NLRP11^−/−^* macrophages (p<0.05 and p<0.01, respectively) and completely abolished in *CASP4^−/−^* and *NLRP11^−/−^ CASP4^−/−^* (p<0.0001) macrophages (Figures 1D, S1B, S1D and S1E). These results further support that, in response to purified LPS, but not bacterial-associated LPS, CASP4 mediates cell death via both NLRP11-dependent and NLRP11-independent pathways.

Pretreatment with the NLRP3 inhibitor MCC950 prior to *S. enterica* LPS electroporation reduced CASP4 and GSDMD processing in THP-1 WT macrophages (p<0.01 and p<0.0001, respectively) (Figures 1D and S1D), indicating functional crosstalk between canonical NLRP3 signaling and CASP4 activation under these conditions. In *NLRP11^−/−^* macrophages, MCC950 treatment reduced GSDMD processing (p<0.001) but did not affect CASP4 processing, demonstrating that NLRP11 is required for optimal activation of both canonical NLRP3 and non-canonical CASP4 inflammasome pathways, consistent with prior observations [9,11,16]. Notably, although MCC950 reduced the amount of active CASP4 and GSDMD under these experimental conditions, the data suggest activation in the presence of MCC950 was nevertheless sufficient to support macrophage cell death following cytosolic LPS delivery (Figure 1B).

Consistent with these results, compared with WT macrophages, release of the IL-1 family cytokines IL-1β and IL-18 from *S. flexneri*-infected or LPS-electroporated *NLRP11^−/−^*, *CASP4^−/−^*, and *NLRP11^−/−^ CASP4^−/−^* macrophages was impaired (Figures S2A-S2F). MCC950 treatment strongly suppressed IL-1β and IL-18 release in all macrophage lines after LPS electroporation, presumably because inhibition of NLRP3 blocks at least part of the downstream signaling required for cytokine maturation and secretion. Collectively, these findings identify NLRP11 as a critical regulator of CASP4-dependent pyroptotic signaling and non-canonical inflammasome activation in response to cytosolic LPS.

### NLRP11 forms an ASC-independent complex with CASP4

The NLRP3 inflammasome functions as a supramolecular organizing center that nucleates caspase 1 activation [20,21]; therefore, we investigated whether NLRP11 similarly assembles a macromolecular complex with CASP4. We found that NLRP11 and CASP4 co-precipitate in a complex with an approximate stoichiometry of 7:1 (CASP4:NLRP11), as determined by densitometry analysis of NLRP11 immunoprecipitation in which the two proteins are expressed at comparable levels (Figures 2A and 2B). Notably, this interaction occurs independently of the adaptor protein ASC, which is essential for canonical NLRP3-CASP1 inflammasome assembly [22,23], as the interaction was observed upon ectopic expression in HEK293T cells, which lack most innate immune proteins, including ASC and CASP4 [16].

**Figure 2.**
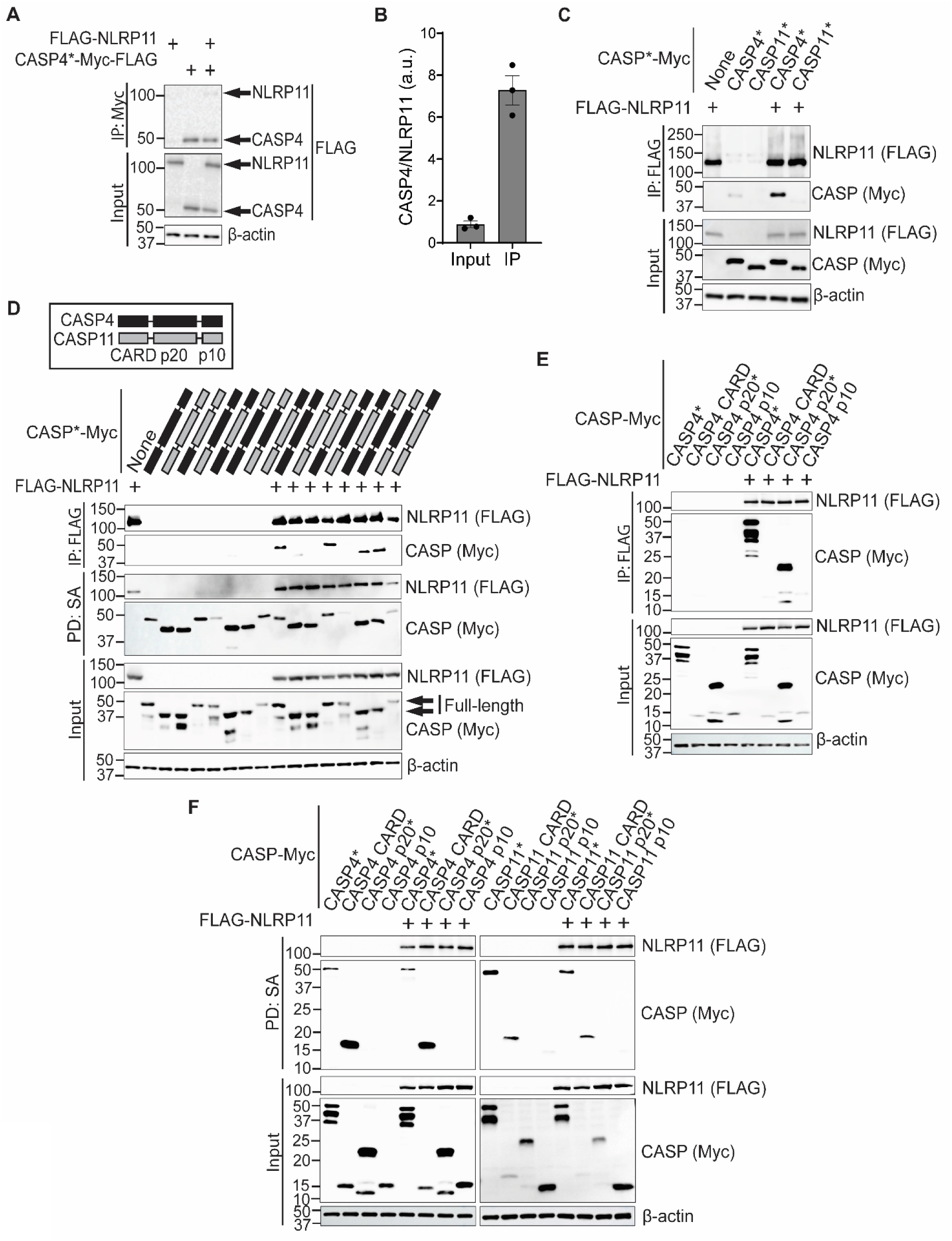
Species-specific interaction of NLRP11 with human CASP4 via its p20 domain. Interaction of NLRP11 and caspases upon heterologous expression in HEK293T cells. (**A**) CASP4 precipitation of human NLRP11. IP: Myc, proteins precipitated with Myc antibody-coated beads. Detection of NLRP11 and CASP4 with anti-FLAG antibody. (**B**) Densitometry of bands corresponding to the experiments represented in **A**. Bars indicate CASP4 signal normalized to NLRP11. a.u., arbitrary units. (**C**) NLRP11 precipitation of human CASP4 or mouse CASP11. (**D**) Precipitation of chimeras of CASP4 and CASP11 by NLRP11 (top 2 blots) and precipitation of indicated caspases and/or NLRP11 by biotin-LPS (3rd and 4th blot). (**E**) Precipitation of domains of CASP4 by NLRP11. (**F**) Precipitation of domains of CASP4 and CASP11 and/or NLRP11 by biotin-LPS. IP: FLAG, precipitation with FLAG antibody-coated beads. PD: SA, precipitation with streptavidin beads. Detection of NLRP11 with anti-FLAG and CASPs with anti-Myc antibodies. CASP*, catalytically inactive caspase. Representative western blots. Molecular weight markers in kDa.

Further supporting ASC independence, immunoprecipitation of NLRP11 by CASP4 in HEK293T cells co-expressing ASC did not result in ASC co-precipitation (Figure S3A). Similarly, in *CASP4^−/−^* or *NLRP11^−/−^ CASP4^−/−^* THP-1 macrophages stably expressing CASP4-Myc, immunoprecipitation using a Myc antibody did not co-precipitate ASC (Figure S3B). In addition, NLRP3 and CASP1 were also not co-immunoprecipitated with CASP4-Myc. Moreover, in THP-1 WT macrophages, immunoprecipitation of CASP4 using a CASP4-specific antibody did not co-precipitate ASC, even upon treatment with extracellular LPS and the NLRP3 agonist nigericin or following LPS electroporation (Figure S3C). These results indicate that ASC does not interact with CASP4 at rest or upon activation of the CASP4 or NLRP3 inflammasome. By contrast, ASC was efficiently co-precipitated with CASP1 upon treatment with extracellular LPS and nigericin, consistent with the known role of ASC in the canonical NLRP3 inflammasome (Figure S3C). Together, these findings are consistent with NLRP11 forming a macromolecular complex with CASP4 in an ASC-independent manner.

### Species-specific interaction of NLRP11 with human CASP4 is via its p20 domain

NLRP11 is present in primates but absent in other mammals [16]. Upon heterologous expression in HEK293T lysates, NLRP11 bound strongly to CASP4 but only weakly to CASP11 (Figure 2C), as we reported previously [16]. Consistent with this, based on a predicted structure of NLRP11, in silico analysis using Pipeline for the Extraction of Predicted Protein-Protein Interactions (PEPPI) [24] predicted an interaction of NLRP11 with CASP4 (log(LR) = 0.053) but not with CASP11 (log(LR) = −0.997).

To identify the CASP4 domain required for interaction with NLRP11, we performed immunoprecipitation of CASP4, CASP11, and CASP4-CASP11 chimeras with NLRP11. We found that the CASP4 p20 domain is necessary for the NLRP11-caspase interaction (Figure 2D, top two blots). The chimeras appeared properly folded, as suggested by their retained ability to bind LPS (Figure 2D, third and fourth blots). By immunoprecipitation with the isolated CASP4 domains (CARD, p20, and p10), we found that the CASP4 p20 domain was sufficient for the interaction with NLRP11 (Figure 2E). As previously described [5], the CARD domain of each CASP4 and CASP11 was sufficient for LPS binding (Figure 2F). Notably, NLRP11 binds LPS regardless of whether CASP4 or CASP11 is present, either as full-length proteins or individual domains. Conversely, CASP4 or CASP11, whether full-length or just their CARD domains, binds LPS independently of NLRP11. Together, these results indicate that NLRP11 specifically interacts with CASP4 via its p20 domain and binds to LPS, suggesting the model that NLRP11 modulates CASP4 engagement of LPS.

### A conserved CASP4 p20 domain residue is required for interaction with NLRP11

We previously observed that CASP4 and CASP1 interact with NLRP11, whereas CASP11 and CASP5 do not [16]. To identify residues within the CASP4 p20 domain required for this interaction, we looked for residues conserved among those caspases that interact with NLRP11 (CASP4 and CASP1) and divergent in those that do not interact with NLRP11 (CASP5 and CASP11). Alignment of CASP4, CASP11, CASP1, and CASP5 revealed three residues in CASP4 (T160, S172, and D174) that meet these criteria (Figure 3A). To assess the contribution of these residues to NLRP11 binding, we generated point mutants in CASP4* (catalytic mutant C258A) in which each residue was substituted with the corresponding amino acid present in CASP11. Immunoprecipitation of CASP4* and its mutants by NLRP11 showed that D174 was required for the interaction, whereas mutations at T160 and S172 had no effect (Figures 3B and 3C). Interaction between NLRP11 and CASP4 D174V was reduced by 65% (p<0.05). Similar results were obtained when NLRP11 was precipitated by CASP4 WT, the catalytic mutant (C258A), or the D174V mutant (Figure S3A). Consistent with these findings, mutational analysis in CASP1 showed that residue D201, analogous to CASP4 D174, was required for CASP1 interaction with NLRP11, whereas CASP1 residues T187 and S199 (analogous to CASP4 T160 and S172) were not (Figure S4).

**Figure 3.**
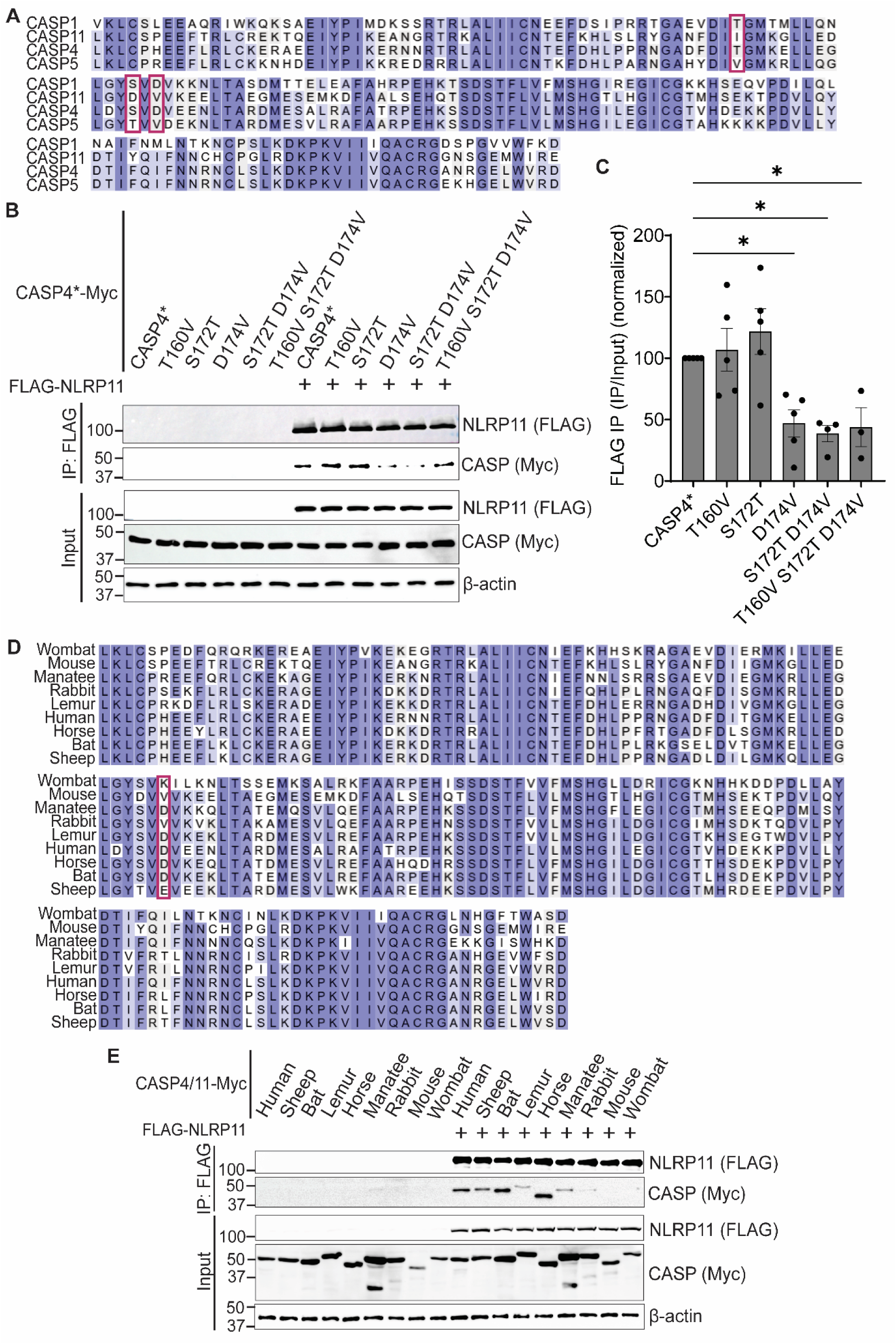
CASP4 residue required for the interaction with NLRP11. (**A**) Alignment of the p20 domains of CASP1, CASP11, CASP4, and CASP5. Three residues conserved between CASP4 and CASP1, but divergent in CASP11 and CASP5, are highlighted. (**B**) Interaction of NLRP11 with indicated CASP4 mutants in HEK293T cells. (**C**) Densitometry of bands corresponding to the experiments represented in panel **B**. Signal normalized to CASP4*. Mean ± SEM. Two-way ANOVA with Tukey’s post hoc test; *, P<0.05. (**D**) Alignment of p20 subunits of CASP4 homologs of non-primate species. The residue corresponding to human CASP4 D174 is highlighted. (**E**) NLRP11 precipitation of CASP4 homologs from indicated species. IP: FLAG, proteins precipitated with FLAG antibody-coated beads. Representative western blots. Detection of NLRP11 with anti-FLAG and CASPs with anti-Myc antibodies. CASP*, catalytically inactive caspase. Molecular weight markers in kDa.

To further assess the functional and evolutionary relevance of CASP4 D174, we aligned CASP4 homologs from multiple non-primate species with human CASP4. D174 was conserved in select non-primate species, including manatee, lemur, horse, and bat (Figure 3D). This conservation correlated with the homologs’ ability to interact with NLRP11; CASP4 homologs retaining an aspartic acid at the position equivalent to human D174 (bat, lemur, horse, and manatee) interacted with NLRP11 (Figure 3E). Notably, the sheep CASP4 homolog also interacted with NLRP11 despite lacking D174; instead, it contains a glutamic acid at this position, which similarly carries a negative charge. In contrast, CASP4 homologs from rabbit and mouse, which contain a valine, and from wombat, which contains a lysine at the equivalent position, failed to interact with NLRP11 (Figures 3D and 3E).

### Multiple NLRP11 domains contribute to CASP4 binding, with the LRR domain mediating the strongest interaction

To determine which NLRP11 domain mediates interaction with CASP4, we examined the ability of full-length NLRP11 and its individual domains (PYRIN, NATCH, and LRR) to precipitate CASP4 p20 domain. The isolated CASP4 p20 interacted with full-length NLRP11 and each individual NLRP11 domain, but the strongest interaction was observed with the LRR domain (p<0.05) (Figures 4A and 4B). To further evaluate these interactions, we immunoprecipitated CASP4/CASP11 chimeras with full-length NLRP11 or its individual domains (Figure 4C). CASP4 interacted with all NLRP11 domains yet displayed the strongest association with the LRR domain. A similar interaction pattern was observed for the CASP11 CARD-CASP4 p20-p10 chimera, supporting a central role for the CASP4 p20 region in NLRP11 binding. In contrast, CASP11 and the CASP4 CARD-CASP11 p20-p10 chimera exhibited only very weak interactions and only with the NLRP11 LRR domain. Together, these results indicate that while multiple NLRP11 domains contribute to CASP4 binding, the LRR domain mediates the strongest interaction, consistent with its established role in ligand and protein recognition [25].

**Figure 4.**
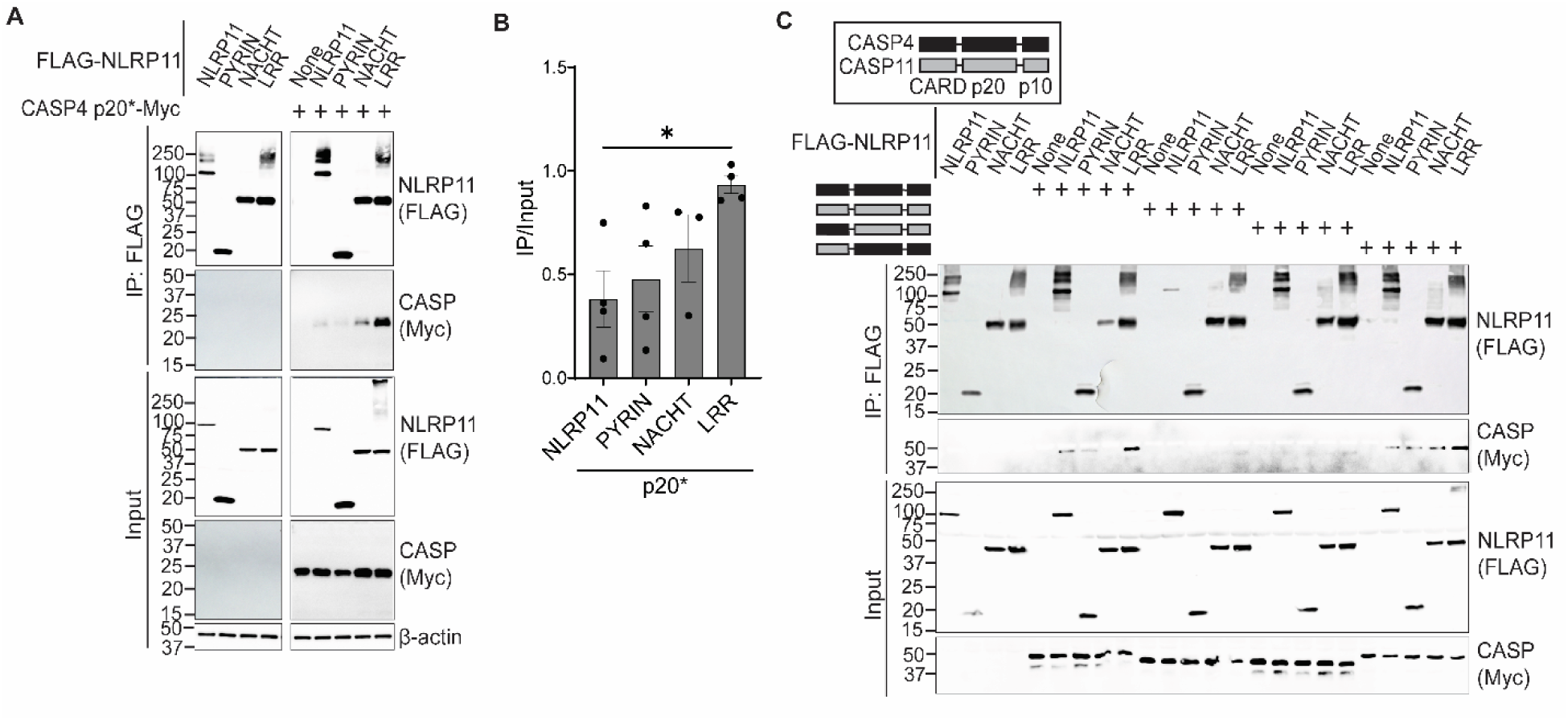
The NLRP11 LRR domain directs interaction with CASP4. (**A**) Precipitation of CASP4 p20* domain by full-length NLRP11 and its individual domains (PYRIN, NACHT, and LRR). (**B**) Densitometry of bands corresponding to the experiments represented in panel **A**. Mean ± SEM. One-way ANOVA with Dunnett’s post hoc test *, P<0.05. (**C**) Precipitation of chimeras of CASP4 and CASP11 by NLRP11 and its individual domains. IP: FLAG, proteins precipitated with FLAG antibody-coated beads. Representative western blots. Detection of NLRP11 and its domains with anti-FLAG and CASPs with anti-Myc antibodies. CASP*, catalytically inactive caspase. Molecular weight markers in kDa.

### NLRP11 and CASP4 interact with LPS independently of each other

To define the CASP4 residues required for LPS binding, we identified those CASP4 residues that align with the CASP11 CARD domain residues previously shown to be required for binding LPS [5]. This analysis identified three sequences within the CASP4 CARD domain, K19, KVR53-55, and QEK62-64, each of which is conserved in CASP11. Point mutations of these residues (K19E, KVR53-55EEA, and QEK62-64EEE) markedly impaired precipitation of CASP4 by LPS (Figure 5A). Combination of the mutations of all three sequences reduced CASP4-LPS binding by 86% (p<0.0001) (Figure 5B). These mutations did not affect the ability of CASP4 to interact with NLRP11 (Figure 5C), indicating that CASP4 binding to NLRP11 and to LPS are mediated by distinct structural determinants. We previously showed that NLRP11 interacts with CASP4 and LPS independently [16]. Consistent with this, biotin-conjugated LPS efficiently precipitated NLRP11 in the absence or presence of CASP4, including in the presence of an LPS-binding-defective CASP4 mutant (Figure 5D). These findings identify residues in the CASP4 CARD domain required for binding NLRP11 and redemonstrate that NLRP11 binds LPS independently of CASP4.

**Figure 5.**
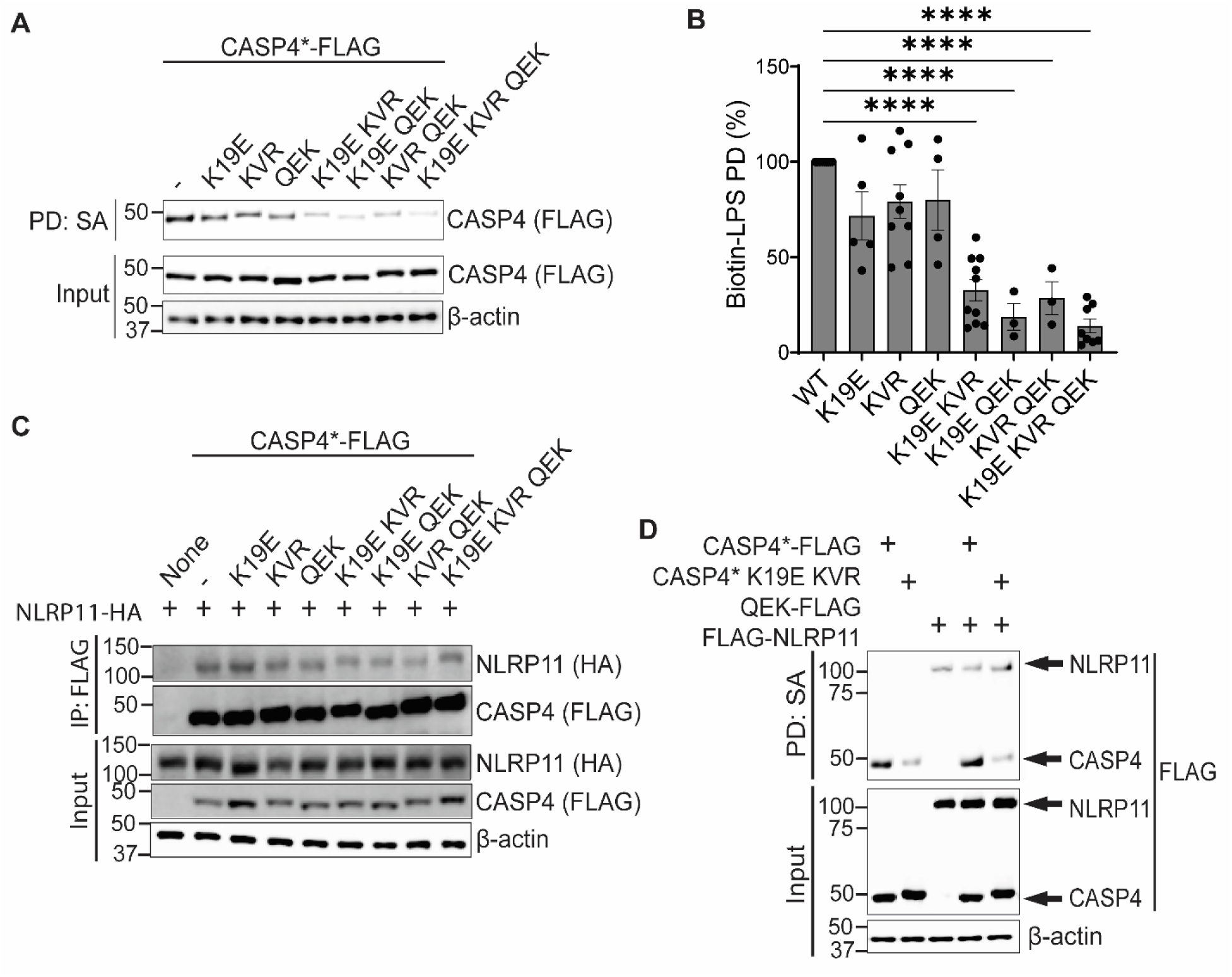
NLRP11 and CASP4 interact independently with LPS. LPS precipitations of lysates of HEK293T cells expressing CASP4 derivatives and/or NLRP11. (**A**) Precipitation of CASP4* or derivatives by biotin-LPS. PD: SA, precipitation with streptavidin beads. (**B**) Densitometry of bands corresponding to experiments represented in panel **A**. Mean ± SEM. Two-way ANOVA with Tukey post hoc test ****, P<0.0001. (**C**) Precipitation of NLRP11 by indicated CASP4 derivatives. IP: FLAG, proteins precipitated with FLAG antibody-coated beads. Detection of NLRP11 with anti-HA and CASPs with anti-FLAG antibodies. (**D**) Precipitation of indicated caspase and/or NLRP11 by biotin-LPS. Representative western blots. CASP*, catalytically inactive caspase. Molecular weight markers in kDa.

### Binding of CASP4 to NLRP11 is essential for CASP4 function

*NLRP11^−/−^*, *CASP4^−/−^*, and *NLRP11^−/−^ CASP4^−/−^* THP-1 macrophages were complemented via stable lentiviral transduction with constructs encoding NLRP11, CASP4 WT, or CASP4 derivatives. All constructs were expressed at both the transcript and protein levels (Figures S5A-S5C). CASP4 WT and derivatives were detected using antibodies to CASP4 or Myc, except for the D174V mutant, which was only recognized by the Myc tag, presumably due to loss of a key epitope caused by the substitution. Levels of NLRP3, CASP1, and GSDMD remained unchanged across all cell lines.

To determine whether interaction with NLRP11 was required for CASP4 function, *CASP4^−/−^* THP-1 macrophages were complemented with a CASP4 mutant defective in NLRP11 binding. Expression of the NLRP11-binding-deficient CASP4 D174V mutant failed to restore WT levels of cell death during *S. flexneri* infection, resulting in a 38% reduction in LDH release compared with CASP4 WT-complemented cells (p<0.05) (Figure 6A). Similarly, following cytosolic LPS delivery by electroporation, CASP4 D174V-expressing macrophages exhibited a 43% reduction in cell death (p<0.0001) (Figure 6B). Consistent with these functional defects, the CASP4 D174V-expressing macrophages showed no detectable CASP4 or GSDMD processing after LPS electroporation (Figure 6C). In each of the complemented THP-1 cell lines, the level of CASP4 was similar, albeit higher than in WT or *NLRP11^−/−^* THP-1 cells (lysates, Figure 6C); the antibody did not detect CASP4 D174V.

**Figure 6.**
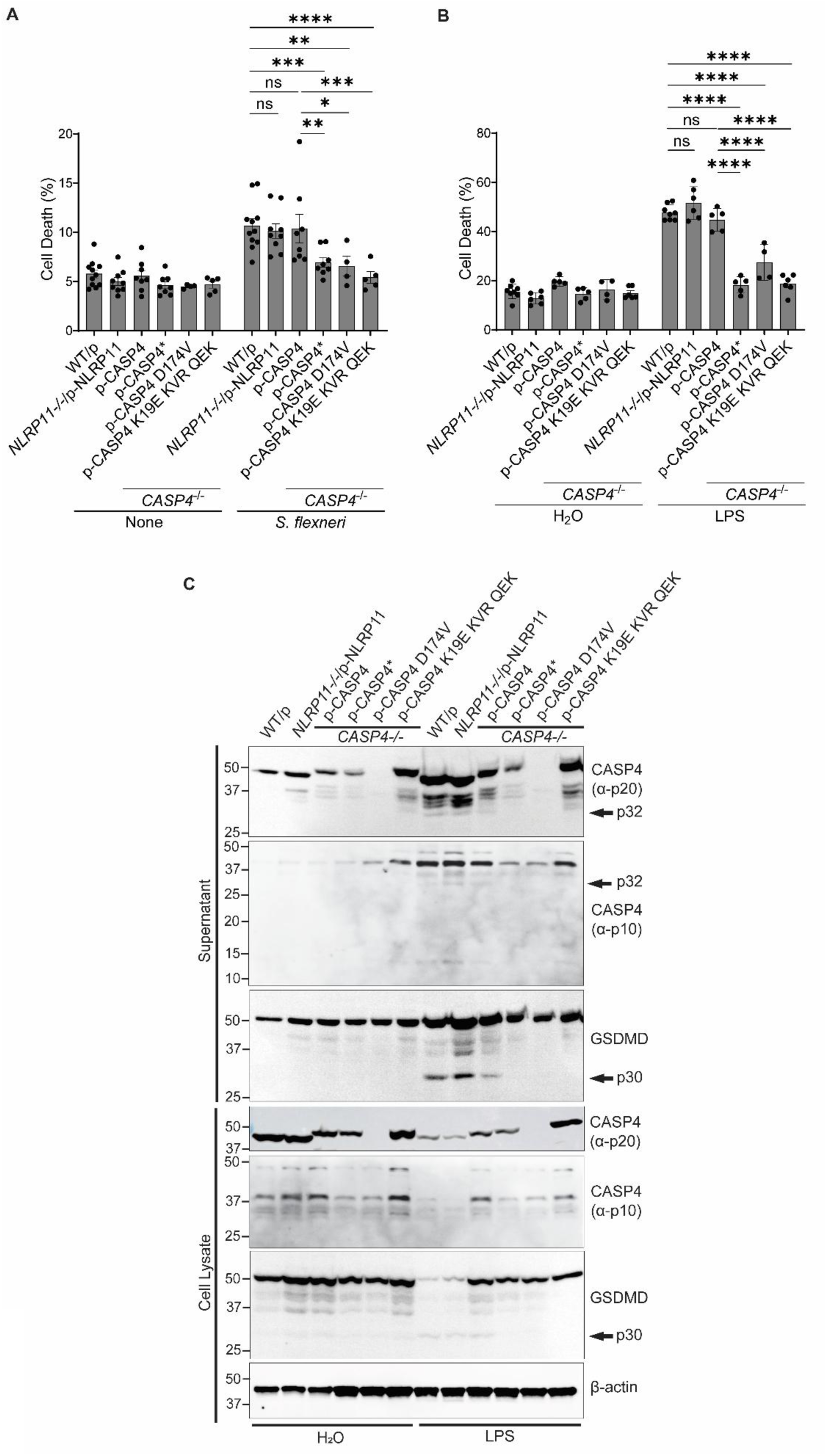
CASP4 interaction with NLRP11 and with LPS are required for efficient macrophage cell death in response to intracellular *S. flexneri* infection or LPS electroporation. (**A-B**). Cell death of THP-1 macrophages upon infection with *S. flexneri* (**A**) or electroporation with *E. coli* O111:B4 LPS (**B**) measured by LDH release, as percentage of Triton X-100-induced cell lysis. Complementation of indicated THP-1 knockouts by transduction with indicated NLRP11 or CASP4 constructs. N ≥ 4 biological replicates. (**C**) Identification of CASP4 residues D174, and K19 KVR53-55 QEK62-64 as required for efficient processing of CASP4 and GSDMD induced by LPS electroporation. Representative western blots. Molecular weight markers in kDa. Mean ± SEM. Two-way ANOVA with Tukey’s post hoc test. ns, not significant; *, P<0.05; **, P<0.01; ***, P<0.001; ****, P<0.0001.

In addition, stable transduction of *NLRP11^−/−^ CASP4^−/−^* THP-1 macrophages with either NLRP11 or CASP4 WT alone failed to restore WT levels of cell death following *S. flexneri* infection or LPS electroporation (Figures S5E and S5F), indicating that both proteins are required for efficient inflammasome activation. In contrast, complementation of *NLRP11^−/−^* macrophages with NLRP11 and of *CASP4^−/−^* macrophages with CASP4 WT restored WT levels of cell death during *S. flexneri* infection (Figure 6A) or following LPS electroporation (Figure 6B), demonstrating efficient rescue of pyroptotic activity; moreover, CASP4 and GSDMD were processed in these two complemented cell lines at levels comparable to WT macrophages after LPS electroporation, indicating that the complemented macrophages behave like WT (Figure 6C). As these experiments were conducted without interferon priming, levels of GBP1 are minimal, indicating that the observed effects of NLRP11 on CASP4 function are independent of GBP1 (Figure S5D). Together, these data demonstrate that both CASP4 and NLRP11 are required and binding of CASP4 to NLRP11 is essential for CASP4 activation and downstream pyroptotic cell death in macrophages.

### CASP4 binding to LPS and catalytic activity are required for non-canonical inflammasome signaling

We next assessed whether CASP4 catalytic activity and LPS binding are also required for CASP4-dependent inflammasome function. Complementation of *CASP4^−/−^* THP-1 macrophages with the LPS-binding mutant CASP4 K19E KVR53-55EEA QEK62-64EEE failed to restore WT levels of cell death during *S. flexneri* infection, resulting in a reductions of 49% compared with CASP4 WT-complemented cells (p<0.001) (Figure 6A). Following LPS electroporation, cell death was reduced by 61% for the LPS-binding mutant compared with CASP4 WT-complemented cells (p<0.0001) (Figure 6B). As expected, complementation with the catalytic mutant CASP4 C258A similarly failed to restore WT levels of cell death during *S. flexneri* infection or following LPS electroporation (Figures 6A-6B).

Consistent with these defects, neither mutant supported detectable processing of CASP4 or GSDMD after LPS electroporation (Figure 6C), indicating that both protease activity and LPS binding are required for non-canonical inflammasome activation. Moreover, release of the IL-1 family cytokines IL-1β and IL-18 from LPS-electroporated THP-1 macrophages was restored to WT levels only in macrophages complemented with NLRP11 and CASP4 WT, but not in cells expressing CASP4 mutants or lacking either protein in the double knockout (Figures S5G and S5H).

Collectively, these findings demonstrate that residues K19, KVR53-55 QEK62-64 are required for CASP4 LPS binding, whereas residue D174 is required for interaction with NLRP11. Together, these interactions are essential for CASP4 processing, GSDMD cleavage, and efficient pyroptotic cell death in response to intracellular *S. flexneri* infection or cytosolic LPS.

### NLRP11 enhances CASP4-dependent LPS binding

To determine whether NLRP11 impacts LPS binding by CASP4, we used a Limulus amebocyte lysate (LAL) assay to quantify binding of purified *E. coli* LPS to CASP4 (WT or LPS-binding-deficient mutants) in the presence or absence of NLRP11. LPS binding to CASP4 in the presence of NLRP11 was increased by 58% (p<0.0001), indicating that NLRP11 promotes LPS binding (Figure 7A). Substitutions of CASP4 residues required for LPS binding (K19 KVR53-55 QEK62-64) markedly impaired LPS binding even in the presence of NLRP11, indicating that CASP4 is the principal ligand for LPS, with NLRP11 promoting LPS binding via an effect on CASP4. Levels of CASP4 and NLRP11 were comparable across all conditions (Figures S6A and S6B).

**Figure 7.**
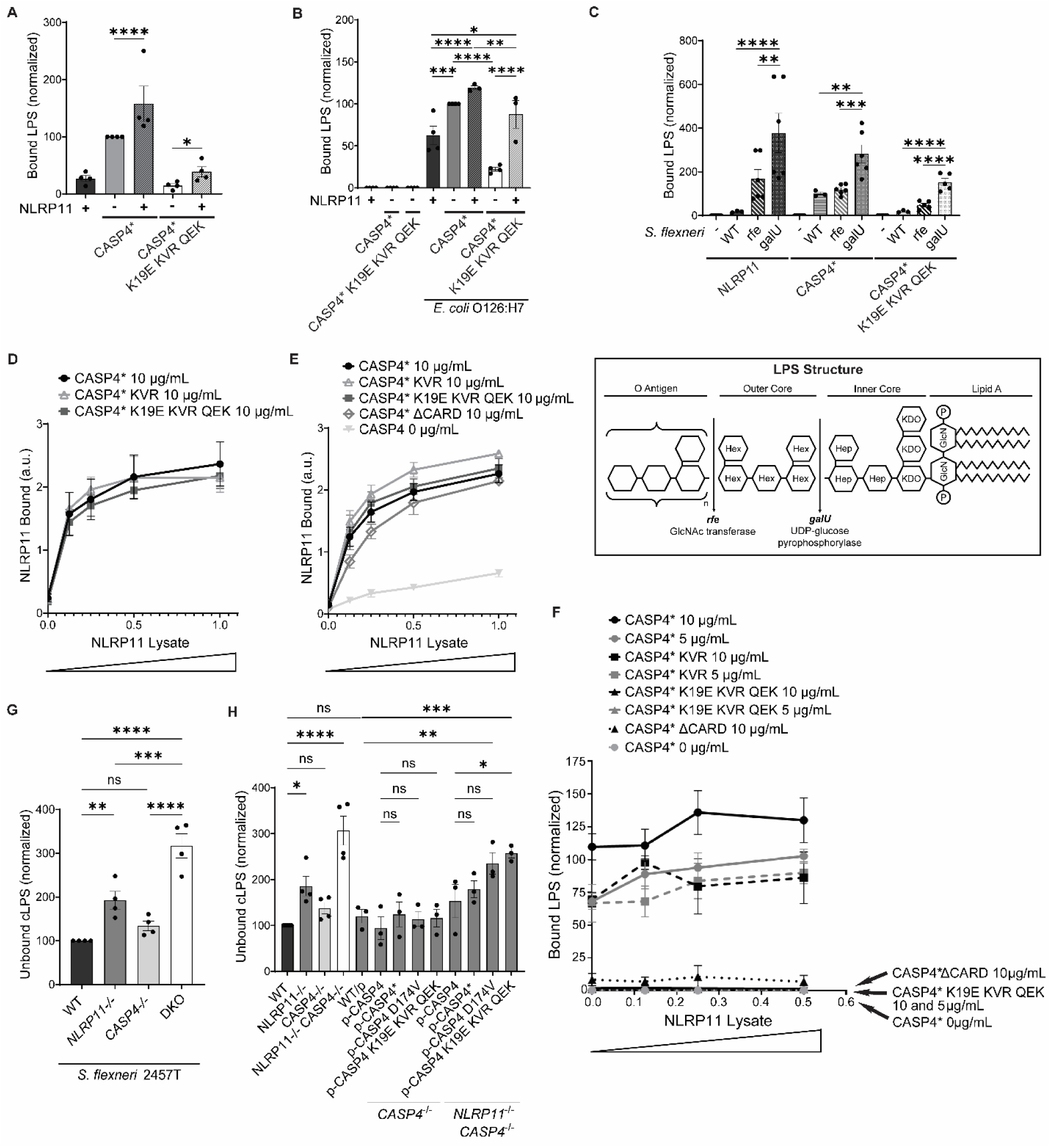
NLRP11 enhances CASP4 binding to LPS in macromolecular complexes and in intact bacteria. (**A-C**). LPS binding to anti-FLAG-coated plates pre-incubated with lysates of HEK293T cells expressing FLAG-NLRP11 and/or CASP4-FLAG. Addition of purified *E. coli* O111:B4 LPS (**A**), intact *E. coli* O126:H7 bacteria (**B**), or intact *S. flexneri* WT, Δ*rfe*, or Δ*galU* bacteria (**C**). Bound LPS was normalized to levels in the CASP4*-alone condition (**A**), CASP4* with *E. coli* O126:H7 (**B**), or CASP4* with WT *S. flexneri* (**C**). Panel **C** also shows the structure of LPS; vertical lines indicate the location of truncations in LPS produced by *S. flexneri* Δ*rfe* and *S. flexneri* Δ*galU*. (**D-E**) Increasing interaction between NLRP11 and CASP4 as a function of concentration of added HEK293T FLAG-NLRP11 lysate, without (**D**) or with (**E**) addition of LPS. FLAG-NLRP11 in transfected HEK293T cells bound to plates coated with purified WT CASP4 or indicated CASP4 derivatives at 10 μg/mL; detection of bound FLAG. N = 3 biological replicates. a.u., arbitrary units. (**F**) LPS binding to plates pre-incubated with lysates of HEK293T cells expressing FLAG-NLRP11 and indicated purified derivatives of CASP4*. (**G-H**) Levels of soluble cytosolic LPS (cLPS) not bound to macromolecular complexes. *S. flexneri*-infected THP-1 macrophages lysed with 0.02% digitonin. Shown is LPS within the separated and filtered cytosolic fractions. Quantified by LAL assay. CASP4*, catalytically inactive caspase. CASP4 K19E KVR QEK, LPS-binding defective caspase. CASP4 KVR, KVR53-55EEA. DKO, *NLRP11^−/−^ CASP4^−/−^*double knockout. Mean ± SEM. Two-way ANOVA with Tukey’s post hoc test; ns, not significant; *, P<0.05; **, P<0.01; ***, P<0.001; ****, P<0.0001.

When intact *E. coli* O126:H7, which expresses a complete O-antigen, was used instead of purified LPS, NLRP11 enhanced LPS binding to CASP4 by approximately 20%, again supporting a role for NLRP11 in facilitating LPS recognition (Figure 7B). In contrast to the binding of purified LPS (Figure 7A), substitutions of CASP4 residues required for LPS binding were associated with only minor decreases in LPS binding when NLRP11 was present, suggesting that NLRP11 may interact with intact bacteria in a manner that is distinct from its interactions with purified LPS. Levels of CASP4 and NLRP11 were comparable across all conditions (Figures S6C and S6D). As GBP1 is absent in HEK293T cells, the observed impact of NLRP11 on CASP4 binding to LPS and intact *E. coli* is independent of GBP1 (Figure S5D). Together, these findings indicate that NLRP11 enhances CASP4 binding to LPS in the cytosol of human macrophages, acting as a primate-specific sensor that facilitates, but does not replace, direct LPS binding by CASP4 during non-canonical inflammasome activation.

Using a series of *S. flexneri* mutants in LPS biosynthesis, we found that the efficiency of LPS binding to both NLRP11 and CASP4 increased as the LPS molecule was truncated, with progressively increased binding to LPS derived from *S. flexneri* WT, Δ*rfe* (lacking O-antigen), and Δ*galU* (lacking outer core and O-antigen). These findings indicate that the distal components of LPS are not required and that lipid A and/or inner core structures are sufficient for binding (Figure 7C). Moreover, they suggest that the distal polysaccharide structures might impose a steric or other impediment to sensor access to the lipid A-proximal regions.

### NLRP11 binds the CASP4 catalytic domain independently of LPS

To define the molecular basis of the NLRP11-CASP4 interaction and to determine whether LPS influences this association, we examined binding between NLRP11 and CASP4 in the presence or absence of LPS. Using CASP4 derivatives purified in a baculovirus system, we observed a concentration-dependent interaction between NLRP11 and CASP4 under both basal conditions and in the presence of LPS (Figures 7D and 7E). Notably, CASP4 ΔCARD bound NLRP11 with efficiency comparable to full-length CASP4, supporting that the CASP4 catalytic domain is sufficient for interaction with NLRP11 and that NLRP11 binding to CASP4 occurs independently of both the CARD domain and LPS, supporting a model in which NLRP11 scaffolds CASP4 through its catalytic domain to promote cytosolic LPS binding. Upon addition of increasing amounts of NLRP11-containing HEK293T cell lysates, increased amounts of LPS bound to purified WT CASP4 (Figure 7F), indicating a dose dependent interaction. In contrast, no LPS binding was detected to CASP4 ΔCARD or the LPS-binding-deficient CASP4 mutant (K19E KVR53-55EEA QEK62-64EEE), even in the presence of NLRP11. These results indicate that enhancement of LPS binding by NLRP11 requires an LPS-binding-competent CASP4, and that NLRP11 alone is insufficient to confer LPS binding under these conditions.

### NLRP11 promotes incorporation of cytosolic LPS into macromolecular complexes

To assess the role of the formation of NLRP11-CASP4 macromolecular complexes on LPS binding, digitonin-fractionation, which selectively extracts soluble cytosolic content, was performed on *S. flexneri*-infected THP-1 macrophages. In the absence of NLRP11 or both NLRP11 and CASP4, elevated levels of cLPS in the soluble fraction (“unbound cLPS”) were detected, with ∼2-fold (p<0.01) and ∼3-fold (p<0.0001) increases, respectively, compared with infected WT THP-1 macrophages (Figure 7G). Bacterial burden and macrophage viability were comparable across all genotypes, indicating that the observed accumulation of the unbound cLPS was not due to differences in bacterial invasion, replication, or cell death (Figures S7A and S7B).

Reconstitution of *CASP4^−/−^* cells with CASP4 WT or CASP4 derivatives was associated with levels of unbound cLPS similar to those observed in WT THP-1 macrophages (Figure 7H). In contrast, reconstitution of *NLRP11^−/−^ CASP4^−/−^* macrophages with CASP4 derivatives defective for binding NLRP11 or LPS resulted in approximately two-fold increases in unbound cLPS (p<0.01 and p<0.001, respectively; Figure 7H). The absence of NLRP11 combined with expression of an LPS-binding-deficient CASP4 resulted in an accumulation of unbound cLPS comparable to that observed in *NLRP11^−/−^CASP4^−/−^*macrophages. As in the parental cell lines, bacterial load and cell viability remained comparable among all complemented conditions, confirming that the observed differences in soluble cLPS reflect defects in inflammasome assembly rather than altered bacterial infection or pyroptotic cell death (Figures S7C and S7D). These findings indicate that NLRP11 enhances CASP4-dependent capture and incorporation of cytosolic LPS into macromolecular complexes.

## Discussion

Cytosolic detection of LPS by inflammatory caspases is a central component of innate immune defense against Gram-negative bacteria. In humans, this pathway is mediated primarily by CASP4, whose activation induces GSDMD cleavage, pyroptotic cell death, and amplification of inflammatory signaling through secondary inflammasome pathways [3–5,7,8]. Although CASP4, and its murine homolog CASP11, can directly bind LPS in vitro, which triggers proximity-dependent oligomerization and activation [5], increasing evidence indicates that efficient non-canonical inflammasome activation in cells depends on spatial organization and accessory host factors that promote ligand capture and signal amplification [12,15,26–28]. For example, the high-mobility group box 1 (HMGB1) protein binds extracellular LPS and facilitates its delivery to the cytosol of mouse macrophages to activate caspase-11 [13,29]. Similarly, host extracellular vesicles provide a spatially organized mechanism for cytosolic delivery of LPS, enabling efficient activation of the non-canonical inflammasome [30]. Notably, inflammasomes and their active components are localized within the cells, and caspase release from the inflammasome is associated with further auto-processing and rapid degradation [31], limiting prolonged or delocalized signaling, emphasizing the importance of spatial and temporal regulation of caspase activity. Here, we identify the primate-specific pattern recognition receptor NLRP11 as a critical upstream determinant of CASP4 activation, enhancing cLPS sensing and pyroptosis through the formation of an ASC-independent complex, with efficient signaling requiring CASP4 enzymatic activity, LPS binding, and direct interaction with NLRP11.

Genetic epistasis analyses suggested that NLRP11 and CASP4 act in the same pathway, as loss of either protein resulted in comparably impaired pyroptotic cell death during *S. flexneri* infection, with no additive defect observed in double-knockout macrophages (Figure 1A), indicating that NLRP11 and CASP4 function within the same signaling pathway rather than in parallel modules. In contrast, cytosolic delivery of purified LPS partially bypassed the requirement for NLRP11 (Figures 1B and S1A), suggesting that NLRP11 is particularly important when LPS is encountered in the context of bacterial infection, a more physiological situation, where ligand availability and spatial organization are more constrained. Consistent with this model, NLRP11 bound intact *E. coli* 2.3-fold more efficiently than purified LPS (Figures 7A and 7B). It is likely that the conformation of bacterially anchored LPS differs from that of purified LPS; we speculate that either NLRP11 has higher affinity for the conformation of LPS that is bacterially anchored, or in addition to binding LPS, NLRP11 binds other molecules on the bacterial surface.

During infection, cLPS does not arise as a uniformly distributed signal but instead originates from localized bacterial lysis or membrane damage, which in interferon primed cells is facilitated by the host antimicrobial guanylate-binding proteins (GBPs) [14,32–35]. The lectin galectin-3 directs GBPs to pathogen-containing vacuoles, enhancing exposure of bacterial LPS to cytosolic sensors in mouse macrophages [36]. In contrast, our study was performed in human macrophages, in which GBPs are minimally expressed in the absence of IFN-γ priming, suggesting that alternative mechanisms are required to support efficient cytosolic LPS sensing. In this context, NLRP11 may function as a primate-specific factor that compensates for limited GBP activity to enable robust CASP4 activation. These data highlight an emerging principle in innate immunity: the physical form and cellular context of microbial ligands can dictate sensor requirements. Accordingly, efficient CASP4 activation likely requires host factors that increase the probability of productive ligand engagement under conditions of limited accessibility or spatial constraint. Our data support a model in which, in unprimed cells, NLRP11 fulfills this role by enhancing the efficiency of CASP4-dependent LPS binding and activation (Figure 7). This context-dependent regulation parallels principles described for other cytosolic pattern recognition pathways, such as cGAS-STING signaling, in which ligand length, localization, and associated host factors determine activation thresholds [37]. By analogy, NLRP11 may function to lower the threshold for CASP4 activation when LPS availability is limiting or spatially constrained.

Mechanistically, we show that NLRP11 interacts with CASP4 via the caspase p20 domain in an ASC-independent manner (Figures 2D, 2E, S3A, and S3B), forming a complex with an approximate stoichiometry of 7:1 (CASP4:NLRP11) (Figures 2A and 2B). Although the stoichiometry may vary under different conditions, our identification of CASP4 residue D174 as essential for NLRP11 binding (Figures 3B and 3C) underscores that this interaction is specific, supporting that spatial coordination by NLRP11 is critical for effective CASP4 activation and downstream pyroptotic signaling. This interaction mode is notable, as most inflammasome scaffolds rely on homotypic CARD-CARD or PYD-PYD interactions to nucleate higher-order assemblies [38,39]. However, as we do not yet know whether the interaction between NLRP11 and CASP4 is direct or indirect, it remains theoretically possible that an adaptor protein that is CARD-like or PYR-like and present in HEK293T cells and macrophages participates in the observed interaction. NLRP11 may facilitate CASP4 oligomerization through the CASP4 CARD domain, effectively functioning as a scaffold that provides sufficient proximity to promote oligomerization and/or positions CASP4 in a signaling-competent state. Supporting this interpretation, recent structural and biophysical studies indicate that non-canonical inflammasomes assemble into heterogeneous, higher-order complexes upon LPS binding, distinct from the canonical filamentous structures [40], which may allow rapid responsiveness to transient cytosolic LPS exposure.

Consistent with our previous work [16], our data indicate that NLRP11 and CASP4 each interact with LPS (Figure 5D). CASP4 recognizes the lipid A and/or the inner core of cLPS through its CARD domain (Figures 5A, 7C and 7F), which drives oligomerization and activation (Figures 6A-6C), confirming previously published data [5]. In intact bacteria, access to lipid A is hindered by the LPS core and O-antigen, and in LPS micelles, lipid Ais buried in the inner assembly. Structural analyses of LPS-protein interactions indicate that distal polysaccharide chains can sterically hinder access to lipid A [41]. Moreover, pathogens such as *S. flexneri* or *Francisella* actively remodel their LPS intracellularly to dampen innate immune recognition and evade inflammasome activation [42,43]. Thus, before CASP4 CARD binding to lipid A transpires, bacterial remodeling or host-mediated processing must occur.

How CASP4 engages lipid A under these conditions is incompletely understood. The published literature and our data provide insights into aspects of this process. In the context of micelles, purified CASP4, including catalytically inactive CASP4 (C258A) and the isolated CARD domain, preferentially binds large LPS micelles and disaggregates them into smaller complexes [44], suggesting a direct role for CASP4 in remodeling LPS assemblies to facilitate signaling. NLRP11 may promote productive CASP4 engagement once lipid A-proximal regions become accessible in the cytosol. Moreover, as NLRP11 appears to interact with intact bacteria (or its LPS) more strongly than with purified LPS (Figure 7B versus 7A), it remains possible that NLRP11 facilitates CASP4 access to lipid A buried in bacterial membranes. Importantly, the mutations in CASP4 CARD that impair LPS binding and inflammasome activation had no effect on the NLRP11-CASP4 interaction (Figure 5C, 7D and 7E). Together, these findings support a model in which NLRP11 functions as an organizing platform that positions CASP4 to efficiently sense and respond to cLPS.

Functional complementation experiments demonstrated that CASP4 activation and non-canonical inflammasome signaling are critically dependent on its interaction with NLRP11, as well as on both its intrinsic enzymatic and LPS-binding functions (Figures 6A-6C, S5G and S5H). A CASP4 mutant defective in NLRP11 binding (D174V) fails to support inflammasome activation despite normal levels of expression (Figures 6A-6C, S5B and S5C), indicating that interaction with NLRP11 is essential for CASP4 function. Similarly, mutants deficient in either catalytic activity (C258A) or LPS binding (K19E KVR53-55EEA QEK62-64EEE) are unable to restore inflammasome activation (Figures 6A-6C, S5B and S5C), confirming and extending previous studies [45,46] showing that enzymatic activity and direct LPS recognition via the CARD domain are requirements for effective non-canonical inflammasome signaling. Collectively, these results are consistent with a model in which NLRP11 scaffolds CASP4, positioning it for productive engagement with cLPS, while CASP4-LPS binding and catalytic activity drive oligomerization, processing, and downstream pyroptotic responses.

We further show that NLRP11 promotes the incorporation of cLPS into CASP4-containing complexes. The absence of NLRP11 resulted in accumulation of unbound cLPS during infection, despite comparable bacterial burdens (Figures 7G-7H and S7A-S7B), indicating a defect in ligand capture rather than ligand availability. Notably, similar phenotypes have been observed upon disruption of host factors that liberate, concentrate, or spatially organize cytosolic LPS, including GBPs and bacteriolytic pathways [33,35].

From an evolutionary perspective, the primate-restricted nature of NLRP11 reflects species-specific differences in innate immune sensitivity. Humans are markedly more sensitive to LPS-induced inflammation and sepsis than mice, a discrepancy that cannot be fully explained by differences in TLR4 signaling alone [17,18,47]. The presence of NLRP11 in primates, but not in lower animals, may represent an adaptation that enhances responsiveness to intracellular Gram-negative bacteria, albeit at the cost of increased inflammatory potential. Consistent with this model, NLRP11 interacts with human CASP4 but not with its murine homolog CASP11 (Figure 2C). Moreover, although CASP4 and CASP5 share overlapping functions in cLPS sensing, NLRP11 selectively binds CASP4 but not CASP5, indicating distinct regulatory mechanisms between these closely related inflammatory caspases [16]. Extending this concept, we observe species-specific differences in the ability of CASP4 homologs from different non-primate mammals to interact with NLRP11 (Figures 3D-3E), suggesting that NLRP11-dependent regulation of non-canonical inflammasome signaling has evolved specifically within the primate lineage.

Taken together, our findings support a model in which cLPS sensing is a multistep process involving ligand release, concentration, capture, and scaffolded caspase activation, with NLRP11 acting at the level of ligand incorporation into signaling-competent CASP4 assemblies. This coordination ensures efficient detection of intracellular bacterial signals and robust inflammatory responses.

## Materials and Methods

### Bacterial strain and growth conditions

*S. flexneri* serotype 2a wildtype strain 2457T [48] was grown overnight in tryptic soy broth at 37°C with aeration. *E. coli* O126:H7 (gift of M. A. Schmidt) was grown overnight in Luria broth at 37°C with aeration.

### Plasmid construction

pCMV-CASP4-FLAG and pCMV6-CASP4-Myc were generated from pCMV-CASP4-Myc-FLAG (gift of S. Shin). pCMV-CASP4-FLAG served as the template to generate the K19E, KVR53-55EEA, QEK62-64EEE, and C258A mutants, and pCMV6-CASP4-Myc was used as the template to generate the T160V, S172T, D174V, and C258A mutants, using QuikChange Site-Directed Mutagenesis (ThermoFisher, EP0501). pcDNA3-NLRP11-HA (gift of H. Wu) was subcloned into pCMV-FLAG to generate pCMV-FLAG-NLRP11. pCMV-FLAG-NLRP11 was used as the template to generate the different NLRP11 subunits (PYRIN, NACHT, and LRR), which were cloned using EcoRI and KpnI. pCMV6-CASP1-Myc was generated by cloning CASP1 into pCMV6-Myc using KpnI and XhoI. The CASP1 mutations (C285A, T187V, S199T, and D201V) were generated using QuikChange Site-Directed Mutagenesis (ThermoFisher, EP0501). pCMV6-CASP11-myc and pCMV6-CASP11*-myc (C254A) were generated by cloning CASP11 in pCMV6-myc using EcoRI and XhoI. The different CASP4 and CASP11 subunits (CARD, p20*, and p10) were cloned into pCMVGG using the Golden Gate protocol [49], with a His6 tag at the N-terminus and a Myc tag at the C-terminus. Combinations of these subunits were cloned into pCMVGG to generate pCMVGG-CASP4/11-Myc chimeras using the same Golden Gate protocol. pCMV-FLAG-ASC was generated by cloning ASC from pET2a-MBP-ASC into pCMV-FLAG using HindIII and EcoRI. pCMV6-CASP4-Myc constructs from different non-primate species (lemur, manatee, horse, sheep, bat, wombat, and rabbit) were cloned from pMSCV2.2 (Addgene, Plasmid #183356) [50] into pCMV6-Myc using KpnI and MluI. Plasmids used for transduction into THP-1 cells (pBABE-FLAG-NLRP11, pBABE-CASP4-Myc, and their mutant derivatives) were generated by cloning the corresponding constructs from pCMV-FLAG-NLRP11 and pCMV6-CASP4-Myc into pBABE using BamHI and SalI. All FLAG tags are 3xFLAG. All C-terminal Myc tags are 1xMyc. DNA used for transfection was prepared LPS-free (Macherey-Nagel, 740410.50).

### Mammalian cell culture

All cell lines were obtained from ATCC. HEK293T cells were cultured in Dulbecco’s Modified Eagle Medium (DMEM; Gibco, 11965118), and THP-1 cells were maintained in RPMI-1640 (Gibco, 21870092). All media were supplemented with 10% (vol/vol) heat-inactivated, sterile-filtered fetal bovine serum (R&D Systems). THP-1 culture medium was additionally supplemented with 10 mM HEPES (Gibco, 15630080) and 2 mM L-glutamine (Gibco, 25030081). Cells were maintained at 37°C in a humidified incubator with 5% CO₂. For differentiation of THP-1 monocytes into macrophage-like cells, 50 ng/mL phorbol-12-myristate-13-acetate (PMA) was added 48 h prior to experiments. When indicated, THP-1 cells were treated with 2 µM of MCC950 (NLRP3 inflammasome inhibitor; InvivoGen, inh-mcc) for 1 h prior to the experiment and maintained in the medium for its duration; and HEK293T and THP-1 cells were treated with 10 ng/mL of IFN-γ (Millipore, Sigma-Aldrich, I17001) for 16 to 20 hours prior to analysis.

### Generation of CRISPR knockouts

THP-1 NLRP11 knockout cells were generated in a previous study [9] using chemically modified sgRNAs (Synthego) electroporated as RNP complexes into freshly cultured THP-1 monocytes with the Neon Transfection System (Invitrogen, MPK5000), following the manufacturer’s instructions. To assemble the editing RNP complexes, 100 pmol of each sgRNA was combined with 25 pmol of SpCas9-2NLS nuclease (Synthego) (4:1 molar ratio) in Resuspension Buffer R (Invitrogen, MPK1069B) and incubated for 15 min at room temperature. THP-1 monocytes were washed in PBS, resuspended in Buffer R, and electroporated with the RNP complexes using two pulses at 1700 V and 2 ms pulse width in the 10 µL Neon tip (1 x 10⁶ cells per reaction). Electroporated cells were immediately transferred into 2.5 mL complete growth medium and plated in a six-well dish. After 72 h at 37°C, 1.5 mL of the culture was moved to a 75-cm² flask, and 1 mL was used for genomic DNA extraction with the DNeasy Blood and Tissue Kit (Qiagen, 69504). Editing efficiency was assessed by PCR, sequencing, and ICE analysis (Synthego). The remaining cells were limiting-diluted into 96-well plates, expanded as single clones, and validated by sequencing to confirm homozygous out-of-frame deletions.

CASP4 knockout THP-1 cells were generated in this study using the same protocol, except that two sgRNAs (Synthego) targeting CASP4 exon 2 (ACAAAAUGUACUGAACUGGA and AGUGAGGAAAUCUUUGCCCA) were delivered together with SpCas9-2NLS nuclease (Synthego) at a 3:1 sgRNA:Cas9 molar ratio.

### Generation of transduced THP-1 cell lines

Retroviral particles were generated by transfecting HEK293T cells with pBABE vectors encoding the indicated inserts (FLAG-NLRP11, CASP4-myc, CASP4 C258A-myc, CASP4 D174V-myc, or CASP4 K19E KVR53–55EEA QEK62–64EEE-myc), together with pAdvantage, pGag-Pol, and pVSV-G. HEK293T cells were seeded at 3 x 10⁶ cells per 100-mm dish in DMEM containing 10% FBS and transfected the following day at 70% confluency. For each dish, a transfection mixture containing FuGENE 6 (Promega, E2691) in Opti-MEM (Gibco, 31985070) was combined with 4.5 µg pBABE, 1.7 µg pAdvantage, 1.7 µg pGag-Pol, and 1.125 µg pVSV-G, incubated for 15 min, and added dropwise to the cells. After 15 h, cells were washed twice in serum-free Opti-DMEM and cultured in DMEM with 20% FBS for 24 h. Viral supernatants were collected 48 h post-transfection, filtered through a 0.45-µm filter, supplemented with 8 µg/mL polybrene, and used immediately. For transduction, THP-1 cells were seeded at 6 x 10⁵ cells per well in 6-well plates, collected the next day, and resuspended in 1.5 mL of viral supernatant, adjusted to 2 mL with RPMI + 10% FBS containing polybrene. Cells were spin-infected for 2 h at 1000 x g at room temperature and selected 48 h later with RPMI + 10% FBS containing 1.5 µg/mL puromycin. Resistant cells were expanded for 5-6 days, cryopreserved, and subsequently clonally isolated by limiting dilution.

### Assessment of transcript levels

To determine levels of NLRP11 and caspase 4 transcripts in transduced THP-1 cells, reverse transcription followed by real-time quantitative PCR (RT-qPCR) was performed using SuperScript IV VILO Master Mix (ThermoFisher Scientific, 11756050), SsoFast EvaGreen Supermix (BioRad, #172-5201), and the QuantStudio™ 5 Real-Time PCR System, 384-well, real-time PCR detection system (Applied Biosystems, A28140) with cycles of 30 s at 95°C, followed by 40 cycles of 5 s at 95°C and 20-30 s at 60°C. Relative expression levels were calculated using the manufacturer’s software for double delta threshold cycle (Ct) method (Design and Analysis Software 2.6.0, Applied Biosystems), using transcripts from housekeeping genes ACTB and GAPDH. Primers used in RT-qPCR were: CASP4-F: GGGATGAAGGAGCTACTTGAGG; CASP4-R: CCAAGAATGTGCTGTCAGAGGAC; NLRP11-F: CTGCTAGTCAAATGAAGAGCCT; NLRP11-R: CAACTTGAGTGTGCGAAGTTTAC; ACTB-F: ATTGCCGACAGGATGCAGAA; ACTB-R: GCTGATCCACATCTGCTGGAA; GAPDH-F: CAACAGCGACACCCACTCCT; and GAPDH-R: CACCCTGTTGCTGTAGCCAAA.

### Bacterial infection

*S. flexneri* infections were performed as previously described [16], using exponential-phase bacteria obtained by back-diluting overnight cultures. THP-1 cells were infected at a multiplicity of infection (MOI) of 10. Bacteria were centrifuged onto host cells at 2000 rpm for 10 min and incubated at 37°C. After 30 min, cells were washed twice and supplemented with 25 µg/mL gentamicin to eliminate extracellular bacteria. For LDH and cytokine measurements, THP-1 cells were seeded in 96-well plates at 5 x 10⁴ cells per well and supernatants were collected at 2.5 h and 5.5 h of infection. For western blot detection of CASP4 and GSDMD, THP-1 cells were seeded in 48-well plates at 5 x 10⁵ cells per well. At the same time points, 100 µL of supernatant was collected and mixed with 35 µL of 4x Laemmli buffer containing cOmplete EDTA-free protease inhibitors (Roche, 5056489001). Whole-cell lysates were prepared in 2x Laemmli buffer. CASP4 and GSDMD processing was quantified by densitometry using ImageJ.

### LPS electroporation

THP-1 cells were electroporated with LPS using the Neon Transfection System (Invitrogen) following the manufacturer’s instructions. Cells were treated with either 2 µg/mL *S. enterica* serovar Minnesota R595 LPS (InvivoGen, tlrl-smlps) or 10 µg/mL *E. coli* O111:B4 LPS (InvivoGen, tlrl-eblps). THP-1 cells were seeded in low-attachment 10-cm dishes with 50 ng/mL PMA in RPMI and differentiated for 48 h, after which they were collected in PBS. For electroporation, 1.5 x 10⁶ cells were pelleted and resuspended in Resuspension Buffer R containing LPS or water (control). Cells were pulsed once using a 10 µL Neon tip at 1400 V and 10 ms pulse width, immediately resuspended in 2 mL RPMI, and plated into 96 and 48-well plates. Cells were incubated for 2.5 h at 37°C before assessing LDH release, CASP4 and GSDMD processing, and secretion of the IL-1 family cytokines.

### LDH assay and cytokine ELISAs

LDH release was measured using the CyQUANT™ LDH and G6PD Cytotoxicity Assays (Invitrogen, C20301) according to the manufacturer’s instructions. Absorbance was recorded at 490 nm and 680 nm immediately after addition of the Stop Solution using a BioTek plate reader. LDH activity was calculated by subtracting the 680 nm background from the 490 nm absorbance, and cytotoxicity for each sample was expressed as a fraction of the total LDH released from cells lysed with the Triton X-100 buffer provided in the kit. Assays of levels of secreted cytokines were performed using the Human Total IL-18 DuoSet ELISA (R&D Biosystems, DY318–05) and ELISA MAX Deluxe Set Human IL-1β (Biolegend, 437004), per the manufacturer’s instructions.

### Precipitation experiments

HEK293T cells were transiently transfected in 10-cm plates using FuGENE 6 (Promega, E2691) in Opti-MEM (Gibco, 31985070). A 10x stock Lysis Buffer (500 mM Tris-HCl, 1.5 M NaCl, 50 mM MgCl₂, pH 7.4) was prepared [32]. Forty-eight hours post-transfection, the medium was removed, and cells were lysed in 575 µL 1x Lysis Buffer containing 0.01% digitonin (Invitrogen, BN2006) and cOmplete EDTA-free protease inhibitors (Roche, 5056489001). Lysates were incubated on ice for 15 min, vortexed for 15 s, and centrifuged at 13,200 rpm for 10 min at 4°C. The resulting supernatants were used for precipitation assays. Lysates from THP-1 cells were prepared in the same way, except that these cells were not transfected. PMA-differentiated THP-1 cells were primed with 1 µg/mL *E. coli* O111:B4 LPS (InvivoGen, tlrl-eblps) for 2.5 h and treated with 10 µM nigericin (InvivoGen, tlrl-nig) for 15 min prior to the assay when indicated. Alternatively, THP-1 cells were electroporated with 10 µg/mL *E. coli* O111:B4 LPS using a 100 µL Neon tip (1400 V, 10 ms pulse width).

For immunoprecipitation, 30 µL of FLAG M2 Magnetic Beads (Millipore-Sigma, M8823) or Pierce Anti-c-Myc Magnetic Beads (ThermoFisher, 88843) were washed twice with Immunoprecipitation Buffer (1x Lysis Buffer containing 0.002% digitonin) and equilibrated in the same buffer at 4°C for 30 min. Beads were blocked with lysate from untransfected cells for 2 h and then incubated with 500 µL of protein lysate for 2 h at 4°C. Beads were washed three times with Immunoprecipitation Buffer and eluted with 50 µL 2x Laemmli buffer.

Streptavidin pulldown was performed as previously described [5] with minor modifications. Lysates were pre-cleared with 30 µL Pierce Streptavidin Magnetic Beads (ThermoFisher, 88816) for 30 min at 4°C to reduce non-specific binding, and then incubated with biotinylated *E. coli* O111:B4 LPS (InvivoGen, tlrl-lpsbiot) loaded onto 30 µL Streptavidin beads for 2 h at 4°C. Beads were washed three times with Immunoprecipitation Buffer and eluted with 50 µL 2x Laemmli buffer.

For precipitation assays using caspase-4 or caspase-1 antibodies in THP-1 cells, the Pierce Classic Magnetic IP/Co-IP Kit (ThermoFisher, 88804) was used according to the manufacturer’s instructions. 10 μg of caspase-4 rabbit monoclonal antibody (Abcam, ab238124) or caspase-1 rabbit monoclonal antibody (Abcam, ab207802) were used. Cells were either primed with LPS and treated with nigericin or electroporated with *E. coli* O111:B4 LPS as described above.

### Purification of recombinant caspase-4

Recombinant human caspase-4 variants, including caspase-4 C258A, caspase-4 C258A KVR53-55EEA, and caspase-4 C258A K19E KVR53-55EEA QEK62-64EEE, were expressed using the Bac-to-Bac Baculovirus Expression System (Invitrogen). Caspase-4 constructs were cloned into pFastBac-HT using BamHI and XhoI. Recombinant bacmids were generated by transforming pFastBac-H-caspase-4 into MAX Efficiency DH10Bac competent cells, followed by blue-white selection on LB agar containing kanamycin, gentamicin, tetracycline, Bluo-gal, and IPTG. White colonies were sub-cultured and screened by PCR using pUC/M13 primers to confirm insertion of caspase-4. Bacmid DNA was purified by isopropanol precipitation, ensuring gentle handling to prevent shearing.

Recombinant baculovirus was produced by transfecting Sf9 insect cells with purified bacmid DNA using Cellfectin II (ThermoFisher, 10362100). Transfected cells were incubated at 27°C for 72 h, and the resulting P1 viral stock was harvested from clarified culture supernatants. P1 virus was amplified to generate P2 and P3 viral stocks by infecting suspension cultures of Sf9 cells at a low multiplicity of infection (MOI) and harvesting supernatants 72 h of infection. Viral stocks were clarified, filtered (0.45 µm), and stored at 4°C protected from light.

For protein expression, Sf9 cells (1 x 10⁶ cells/mL) were infected with P3 viral stock at 0.5-1% (v/v) and cultured for 72 h at 27°C. Cells were harvested by centrifugation, and pellets were stored at −20°C until purification. All purification steps were performed on ice or at 4°C using pre-chilled buffers. Cell pellets were resuspended in lysis buffer (25 mM HEPES, pH 7.4, 150 mM NaCl, 10 mM imidazole) supplemented with 1 mM PMSF and 1 mM benzamidine, lysed by sonication, and then clarified by centrifugation. The soluble fraction was applied twice to Ni-NTA agarose resin (Qiagen, 154033938) in a gravity flow column (Biorad Econo-Pac Chromatography Colums, 7321010) pre-equilibrated in lysis buffer. The resin was washed with high-salt wash buffer (25 mM HEPES pH 7.4, 400 mM NaCl, 25 mM imidazole), and caspase-4 was eluted in elution buffer (25 mM HEPES pH 7.4, 150 mM NaCl, 250 mM imidazole).

Eluted protein was concentrated to ∼0.5 mL using a 10-kDa MWCO centrifugal concentrator (Amicon Ultra 15 ml, Millipore, UFC901024) and further purified by size-exclusion chromatography on a Superdex 200 Increase 10/300 GL column (Sigma, GL 28-9909-44) equilibrated in SEC buffer (25 mM HEPES pH 7.4, 150 mM NaCl). Fractions containing caspase-4, identified by SDS-PAGE and Coomassie staining, were pooled, supplemented with 10% glycerol, snap-frozen in liquid nitrogen, and stored at −80°C.

Recombinant human caspase-4 lacking the CARD domain (ΔCARD) was purified from *E. coli* as previously described [51].

### Plate binding assays

Clear, flat-bottom Nunc MaxiSorp ELISA plates (Thermo Fisher, 442404) were coated overnight at 4 °C with the indicated concentrations of purified recombinant caspase-4 proteins, including caspase-4 C258A, caspase-4 lacking the CARD domain (ΔCARD), caspase-4 C258A KVR53–55EEA, and caspase-4 C258A K19E KVR53–55EEA QEK62–64EEE, diluted in 100 mM carbonate–bicarbonate coating buffer (28.6 mM Na₂CO₃, 71.4 mM NaHCO₃, pH 9.6) (Fig. 6D-F). For experiments shown in Figures 6A-6B, anti-FLAG-coated plates were used. Plates were washed three times with PBS and blocked for 2 h at room temperature with 1% bovine serum albumin (BSA; Fisher, BP9700100) in PBS.

HEK293T cells transiently expressing FLAG-NLRP11, CASP4 C258A-myc and derivatives, or untransfected control cells were lysed in lysis buffer (50 mM Tris-HCl pH 7.4, 150 mM NaCl, 5 mM MgCl₂) supplemented with 0.01% digitonin (Invitrogen, BN2006) and cOmplete EDTA-free protease inhibitors (Roche, 5056489001). Lysates were serially diluted in untransfected HEK293T cell lysate, and the indicated concentrations were incubated on pre-blocked plates for 1 h at room temperature. Plates were washed three times with PBS to remove unbound material.

Where indicated, purified *E. coli* O111:B4 LPS (5 µg/mL in PBS containing 1% Tween-20), intact *E. coli* O126:H7, or *S. flexneri* 2457T wildtype (WT), *rfe*, or *galU* mutant strains were added and incubated for 1 h at 37 °C. For detection of FLAG-NLRP11, wells were incubated with 0.8 µg/mL rabbit anti-FLAG antibody (Sigma, F7425-.2MG) for 1 h at room temperature, followed by washing and incubation with 0.16 µg/mL HRP-conjugated goat anti-rabbit secondary antibody (Jackson, 111-035-144) for 1 h. To detect plate-bound caspase-4, wells were incubated with 0.5 µg/mL anti-caspase-4 antibody (Santa Cruz, sc-56056), followed by washing and incubation with 0.16 µg/mL HRP-conjugated goat anti-mouse secondary antibody (Jackson, 115-035-003), each for 1 h.

After six washes with PBS, 50 µL of One Step Ultra-TMB substrate (Fisher Scientific, PI34029) was added to each well and incubated for 5 min. Reactions were stopped with 4 N sulfuric acid, and absorbance was measured using a BioTek plate reader. To quantify LPS bound to the plates, LPS levels were measured using a chromogenic Limulus amebocyte lysate (LAL) assay according to the manufacturer’s instructions (Associates of Cape Cod, #E0005 and #C0031).

### Cytosolic LPS assay

This protocol was adapted with modifications from a previously described method [52]. Briefly, THP-1 cells were seeded in 6-well plates at a density of 2.5 x 10⁶ cells per well and treated with 50 ng/mL PMA. Two days later, cells were infected with *S. flexneri* at an MOI of 10, as described above. Two hours of infection, cells were washed three times with pre - warmed PBS and then treated with 500 µL pre-warmed Trypsin-EDTA (0.25%, Gibco, 25200056). After a 5-min incubation at 37°C, cells were resuspended in 500 µL pre-warmed RPMI containing 10% FBS and transferred to 1.5-mL Eppendorf tubes, which were spun at 100 x g for 10 min. Supernatants were discarded, and pellets were washed twice with PBS before being resuspended in 500 µL ice-cold 0.02% digitonin in PBS. Lysed cells were centrifuged at 400 x g for 10 min to separate soluble (cytosolic) and residual fractions. Cytosolic fractions were diluted 1:2 in endotoxin-free water and filtered (0.2 µm) to remove intact bacteria. Filtered cytosolic fractions were diluted 1:10, after which LPS was quantified using a chromogenic LAL assay according to the manufacturer’s instructions (Associates of Cape Cod, #E0005 and #C0031). Residual pellets were resuspended in 500 µL of 0.5% Triton X-100 and plated to quantify bacterial infection. LDH assays were performed in parallel to assess cell death. Cytosolic LPS levels were normalized to the number of cells per condition.

### Antibodies

Caspase-4 (Santa Cruz, sc-56056), gasdermin D (Santa Cruz, sc-81868), and ASC (Santa Cruz, sc-514414) mouse monoclonal antibodies were used at 0.5 μg/mL (1:200). Caspase-4 rabbit polyclonal antibody (BioRad, AHP964) was used at 1 μg/mL (1:1000). FLAG M2 mouse monoclonal antibody (Sigma, F3165) was used at 76 ng/mL (1:5000). Caspase-1 rabbit monoclonal antibody (Abcam, ab207802) was used at 0.5 μg/mL (1:1000). Myc rabbit monoclonal antibody (Cell Signaling, 2278) was used at 1:1000. NLRP3 mouse monoclonal antibody (Adipogene, AG-20B-0014-C100) was used at 1 μg/mL (1:1000). HA mouse monoclonal antibody (Biolegend, 901501) was used at 0.5 μg/mL (1:2000).

For western blots of FLAG or Myc immunoprecipitations, the secondary antibody used was TrueBlot horseradish peroxidase-conjugated anti-mouse Ig (Rockland, 18-8817-31) at 1:1000. For caspase-4 or caspase-1 immunoprecipitations, the secondary antibody used was Clean-Blot IP Detection Reagent (ThermoFisher, 21230) at 1:1000. For detecting β-actin, horseradish peroxidase-conjugated anti-β-actin antibody (Sigma, A3854) was used at 1:10000. For all other western blots, the secondary antibody used was horseradish peroxidase-conjugated goat anti-rabbit or goat anti-mouse IgG antibodies (Jackson ImmunoResearch, 115-035-003 or 111-035-144). All antibodies were diluted in 5% milk in Phosphate-Buffered Saline with 0.1% Tween (PBST).

### Statistics and reproducibility

Unless otherwise indicated, all results represent data from at least three independent biological replicates. In bar graphs, each symbol corresponds to an individual biological replicate. Statistical analyses were performed using GraphPad Prism version 10.

## Acknowledgements

We thank M. A. Schmidt, S. Shin, and H. Wu for reagents used in this study. This work was supported by NIH R01AI173030 (to M.B.G.), NIH T32AI007061 (to M.L.G.M.), and a Massachusetts General Hospital Executive Committee on Research Fund for Medical Discovery Fundamental Research Fellowship Award GR0244866 (to M.L.G.M.).

## Notes

### Competing Interest Statement

The authors have declared no competing interest.

