## Supplemental Material for "NLRP11 promotes non-canonical inflammasome activation in human macrophages by enhancing caspase-4 recognition of cytosolic lipopolysaccharide"

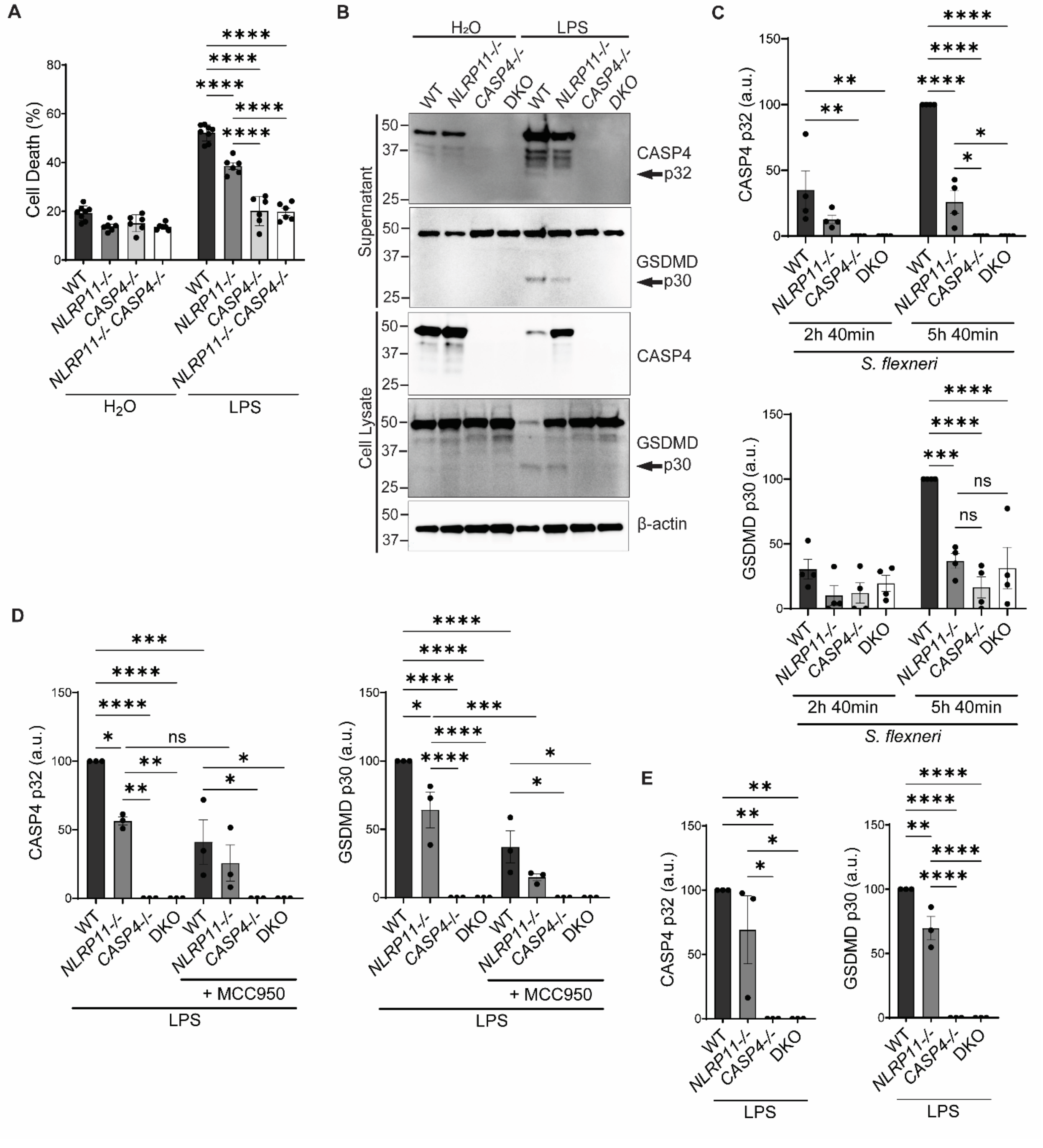


**Supplemental Figure S1. NLRP11 is required for CASP4 and GSDMD processing induced by *S. flexneri* infection or LPS electroporation.** (**A**) Cell death of THP-1 macrophages upon electroporation with *E. coli* O111:B4 LPS, measured by LDH release, as percentage of Triton X-100-induced cell lysis. N ≥ 4 biological replicates. (**B**) NLRP11 is required for the processing of

CASP4 and gasdermin D (GSDMD), and CASP4 is required for GSDMD processing induced by *E. coli* LPS electroporation. Processing of CASP4 yields a p32 and/or p20 polypeptide, and processing of GSDMD yields a p30 fragment. (**C**-**D**) Densitometry of processed bands corresponding to experiments represented in **Figures 1C** and **1D**. (**E**) Densitometry of processed bands corresponding to experiments represented in panel **B** of this figure. Representative western blots. Densitometry values are normalized to WT *S. flexneri* infection at 5 h 40 min or to WT LPS electroporation. Molecular weight markers in kDa. a.u., arbitrary units. DKO, *NLRP11*$^{-/-}$ *CASP4*$^{-/-}$ double knockout. Mean ± SEM. Two-way ANOVA with Tukey's post hoc test; ns, not significant; *, P<0.05; **, P<0.01; ***, P<0.001; ****, P<0.0001.

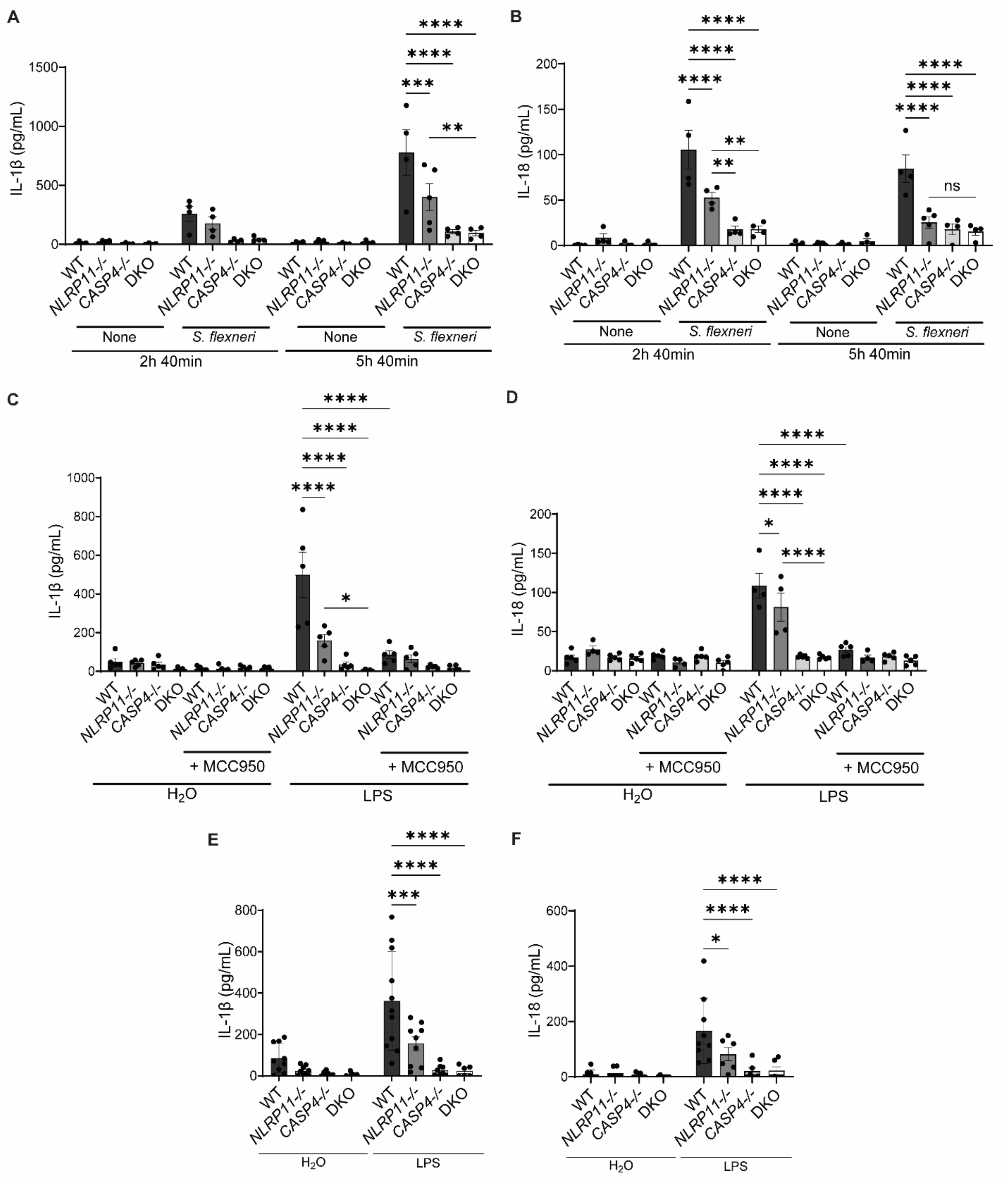


**Supplemental Figure S2. Dependence on NLRP11 and CASP4 of release of IL-1 cytokines from THP-1 macrophages upon *S. flexneri* infection or LPS electroporation.** Release of IL-1β, and IL-18 from WT, *NLRP11*$^{-/-}$, *CASP4*$^{-/-}$ and *NLRP11*$^{-/-}$ *CASP4*$^{-/-}$ double knockout (DKO) THP-1 macrophages. IL-1β (**A, C,** and **E**), and IL-18 (**B, D,** and **F**) levels in the supernatants of macrophages infected with *S. flexneri* for the indicated times (**A**-**B**), electroporated with *S. enterica* serovar Minnesota LPS (**C**-**D**), or electroporated with *E. coli* O111:B4 LPS (**E**-**F**). N ≥ 4 biological replicates. Mean ± SEM. Two-way ANOVA with Tukey's post hoc test; *, P<0.05; **, P<0.01; ***, P<0.001; ****, P<0.0001.

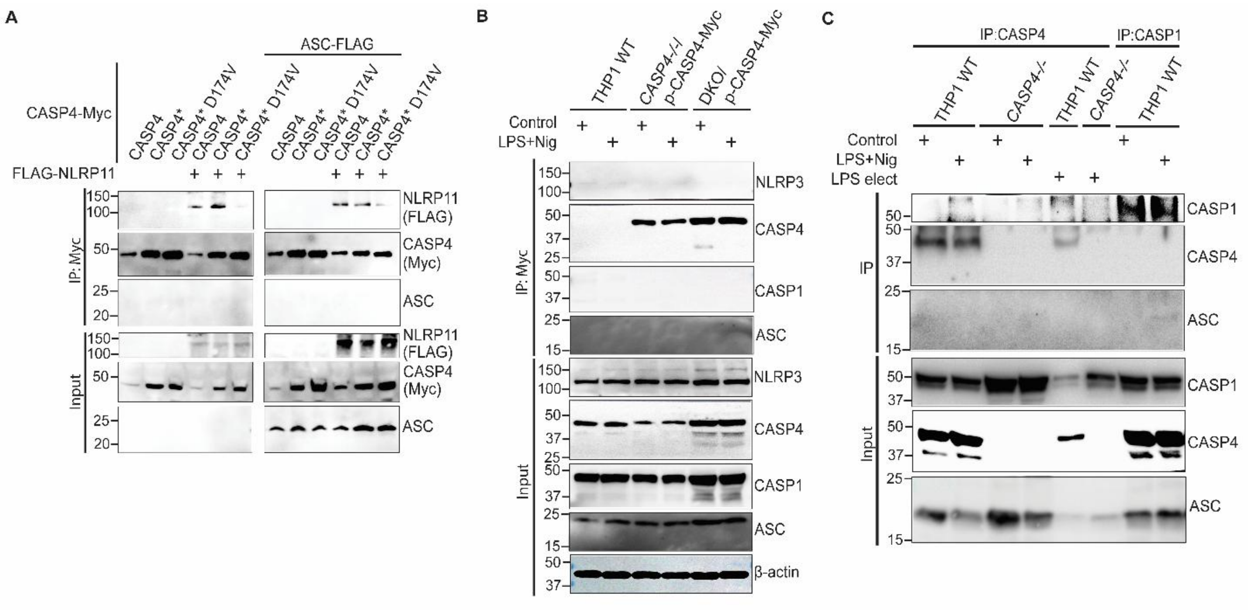


**Supplemental Figure S3. Interaction between NLRP11 and CASP4 does not require the adaptor ASC.** (**A**) CASP4 precipitation of NLRP11, but not ASC, upon heterologous expression in HEK293T cells. IP: Myc, proteins precipitated with Myc antibody-coated beads. Detection of NLRP11 with anti-FLAG and CASPs with anti-Myc antibodies. CASP4*, catalytically inactive caspase. (**B**) CASP4 precipitation from THP-1 macrophage lysates, untreated (control) or treated with extracellular LPS and nigericin. No co-precipitation of NLRP3, CASP1, or ASC was detected. Indicated THP-1 macrophages. DKO, *NLRP11*$^{-/-}$ *CASP4*$^{-/-}$ double knockout. IP: Myc, proteins precipitated with Myc antibody-coated beads. (**C**) Precipitation of CASP4 or CASP1 in THP-1 macrophage lysates, untreated (control) or treated with extracellular LPS and nigericin, or upon LPS electroporation. After treatment with extracellular LPS and nigericin, CASP1 but not CASP4 precipitates ASC. IP: Protein A/G magnetic beads coated with anti-CASP4 or anti-CASP1 antibodies. Representative western blots. Molecular weight markers in kDa.

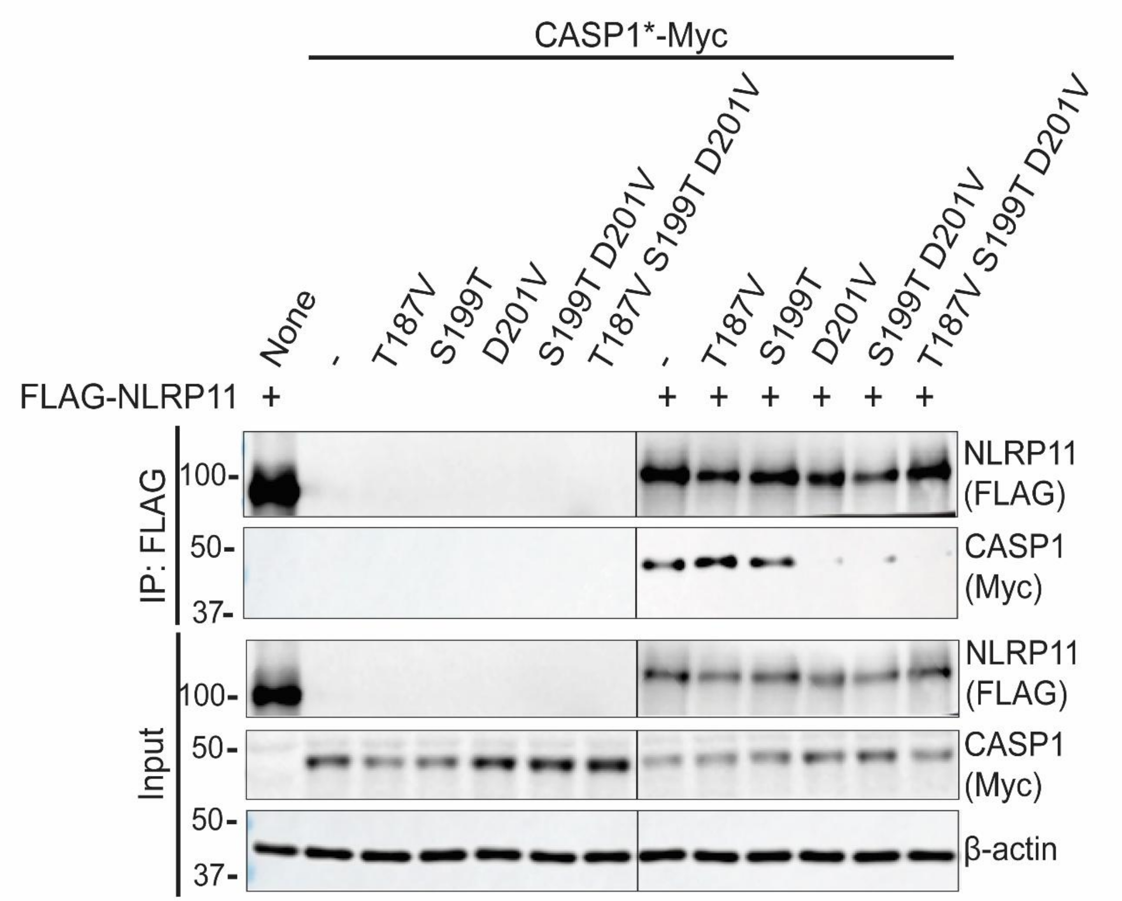


**Supplemental Figure S4. Identification of the CASP1 residue required for interaction with NLRP11.** NLRP11 precipitation of CASP1 mutants from HEK293T lysates. IP: FLAG, proteins precipitated with FLAG antibody-coated beads. Representative western blot. Detection of NLRP11 with anti-FLAG and CASP1 with anti-Myc antibodies. CASP1*, catalytically inactive caspase. Molecular weight markers in kDa.

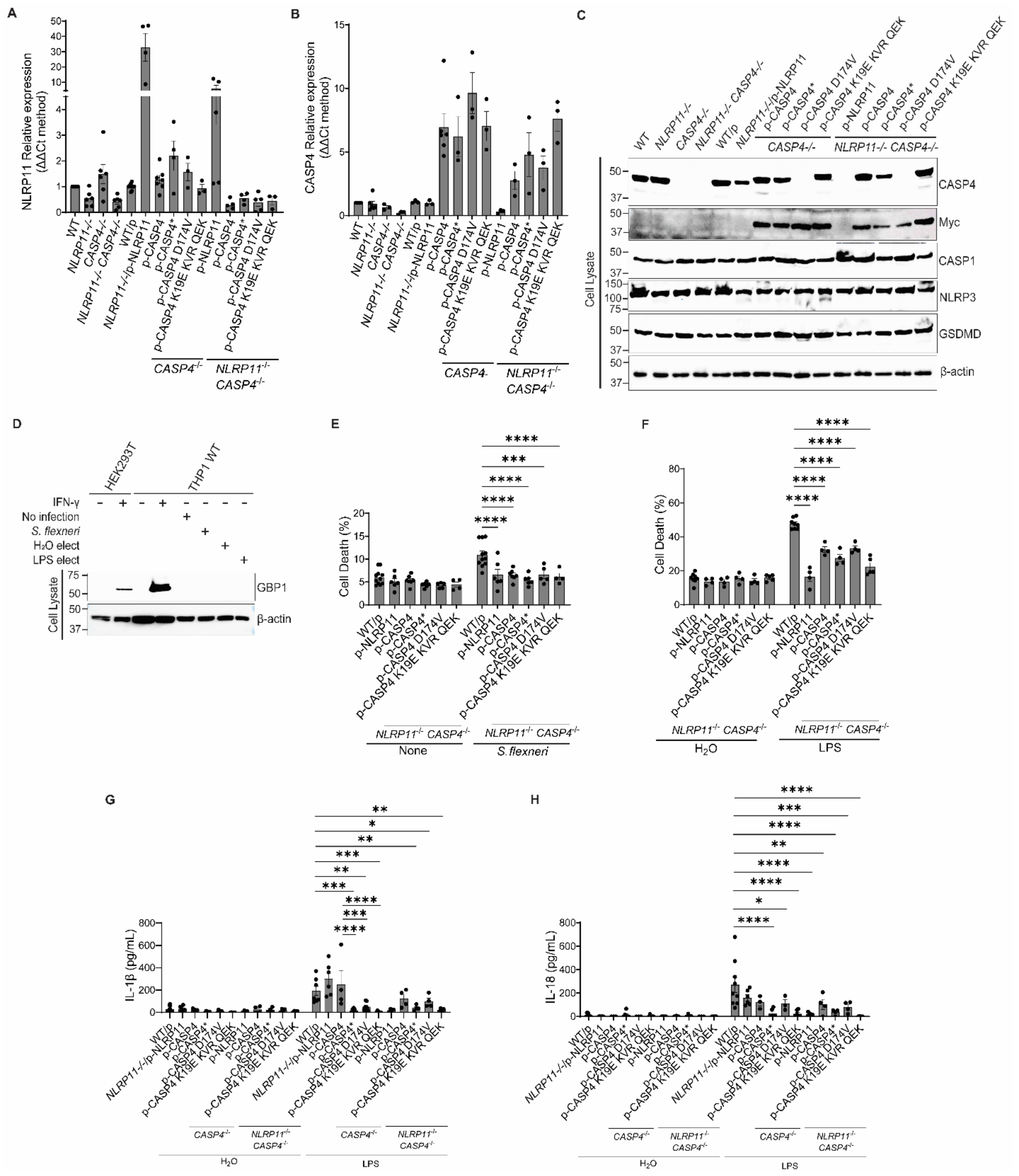


**Supplemental Figure S5. Characterization of THP-1 knockout and complemented cell lines.** (**A-B**). RT-qPCR analysis of NLRP11 (**A**) or CASP4 (**B**) expression in the THP-1 cell lines generated in this study. GAPDH and ACTB, housekeeping genes. (**C**) Levels of CASP4, CASP1, NLRP3, and GSDMD in the indicated THP-1 cell lines. Representative western blots. CASP4 constructs are Myc-tagged. CASP4 D174V is not recognized by the CASP4 antibody. Molecular weight markers in kDa. (**D**) Levels of GBP1 in the indicated cell lines. Representative western blots. Molecular weight markers in kDa. (**E-F**) Cell death of indicated THP-1 macrophages upon infection with *S. flexneri* (**E**) or electroporation with *E. coli* O111:B4 LPS (**F**), measured by LDH release, as percentage of Triton X-100-induced cell lysis. N ≥ 4 biological replicates. (**G-H**). NLRP11 and CASP4-dependence of release of IL-1 cytokines from indicated THP-1 macrophages upon electroporation of *E. coli* O111:B4 LPS. Levels of IL-1β (**G**), or IL-18 (**H**) in the cell supernatants. N ≥ 4 biological replicates. DKO, $NLRP11^{-/-}$ $CASP4^{-/-}$ double knockout. Mean ± SEM. Two-way ANOVA with Tukey's post hoc test; *, P<0.05; **, P<0.01; ***, P<0.001; ****, P<0.0001.

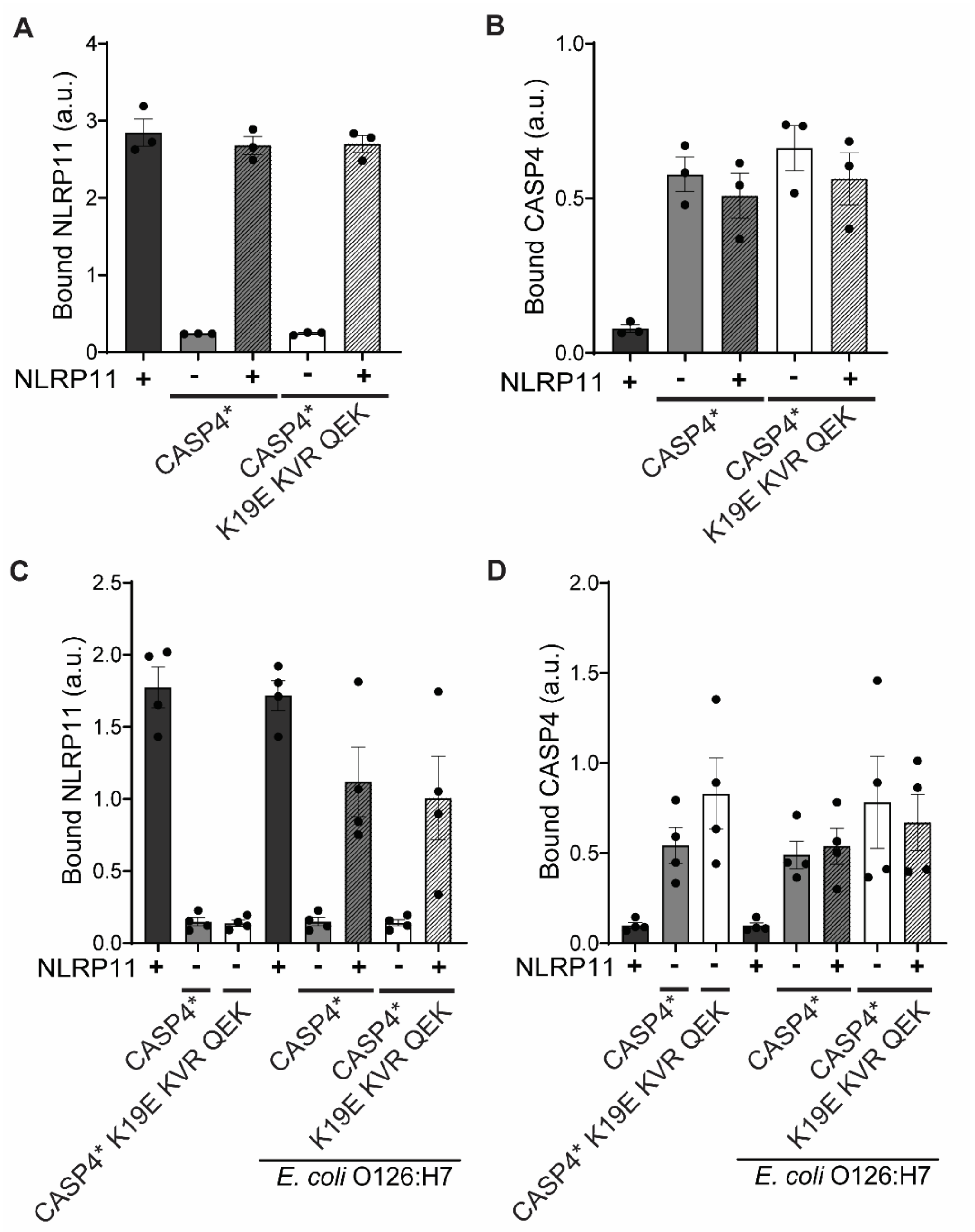


**Supplemental Figure S6. Levels of bound NLRP11 and CASP4.** ELISA detection of levels of NLRP11 or CASP4 constructs bound to anti-FLAG-coated plates in the experiments shown in **Figures 7A** and **7B**. Addition of LPS (**A-B**) or intact *E. coli* O126:H7 (**C-D**). CASP4*, catalytically inactive caspase. CASP4 K19E KVR QEK, LPS-binding defective caspase. a.u., arbitrary units.

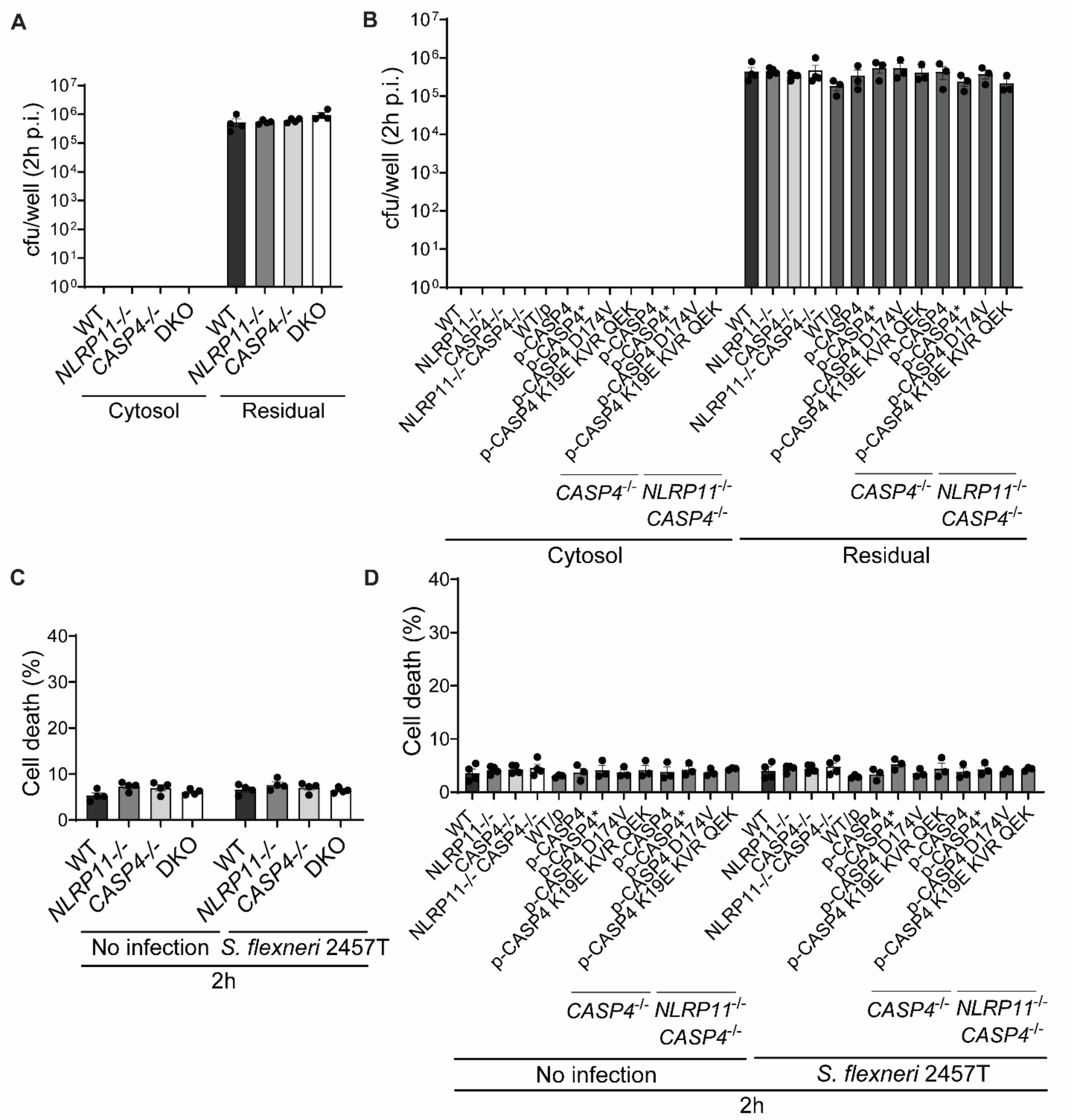


**Supplemental Figure S7. Accumulation of unbound cytosolic LPS is independent of bacterial burden and macrophage viability.** (**A-B**) Quantification of bacteria within the cytosolic and residual fractions (cfu/well) corresponding to the experiments shown in **Figures 7G** and **7H**. (**C-D**) Lack of increase in cell death, measured by LDH release at 2h of infection, corresponding to the experiments shown in **Figures 7G** and **7H**. CASP4*, catalytically inactive caspase. CASP4 K19E KVR QEK, LPS-binding defective caspase. DKO, *NLRP11*$^{-/-}$ *CASP4*$^{-/-}$ double knockout. p.i., post-infection. Mean ± SEM. Two-way ANOVA with Tukey's post hoc test.
